# Enforced MYC expression selectively redirects transcriptional programs during human plasma cell differentiation

**DOI:** 10.1101/2024.04.18.589889

**Authors:** Panagiota Vardaka, Eden Page, Matthew A Care, Sophie Stephenson, Ben Kemp, Michelle Umpierrez, Eleanor O’Callaghan, Adam Mabbutt, Roger Owen, Daniel J Hodson, Gina M Doody, Reuben M Tooze

## Abstract

MYC provides a rheostat linking cell growth and division during plasma cell (PC) differentiation. Precise control of MYC is central to the network controlling differentiation. Deregulation of MYC drives transformation in aggressive B-cell neoplasms and is often accompanied by apoptotic protection conferred by BCL2. We assess how MYC and BCL2 deregulation impacts on the ability of human B-cells to complete PC differentiation. Under permissive conditions for PC differentiation we find such deregulation does not transform cells. While driving loss of normal PC surface phenotype, MYC deregulation has little impact on components of regulatory circuitry controlling B-cell identity. This contrasts with profound impact on initiation of secretory output and secretory reprogramming, coupled to dampening of *XBP1* and immunoglobulin gene enhancement and a shift toward distinct metabolic programs. The establishment of this aberrant state depends on MYC homology boxes (MB0 and MBII). Dependence on MBII is profound and resolves to residue W135.

## Introduction

The transcription factor c-MYC (MYC) was the first oncogene deregulated by chromosomal translocation to be identified in B-cell lymphoma.[1–3] It is one of the most frequently deregulated oncogenes in aggressive lymphoid cancer but also acts as a central regulator of physiological lymphocyte growth and proliferation.[4–9] MYC acts as a sequence specific transcription factor of the basic-helix-loop-helix (bHLH) domain family, occupying E-box DNA sequence elements in complex with its obligatory partner MAX.[10–13] With high MYC expression its genomic occupancy spreads to sites with non-consensus E-box motifs, and MYC occupancy correlates with RNA polymerase loading at primed promoters and with release of paused polymerases.[11, 12, 14–17] Distinct models of MYC function support action as a global enhancer of prevailing active promoters, and as a more selective regulator of specific gene programs that overlap between multiple cell types.[11, 15–18] Exit from cell cycle and cellular terminal differentiation is linked to repression of MYC and nuclear exclusion.[19] In B-cell differentiation to the plasma cell (PC) stage repression of *MYC* has been attributed to the transcriptional factor BLIMP1/PRDM1.[20–23]

MYC-driven cellular transformation depends on DNA binding and on the MYC N-terminal transactivation domain (TAD).[13, 24, 25] This TAD contains evolutionarily conserved regions, the MYC boxes (MB), that are responsible for distinct co-factor interactions.[18, 26, 27] MBI contains a phosphodegron sequence controlling MYC degradation via the proteasome.[28–30] A crucial residue in MBI is T58 which is frequently mutated in aggressive lymphoma.[30–32] MB0 and MBII are implicated in transactivation and transformation activity with MBII identified as essential for MYC driven transformation.[33–35] MBII mediates recruitment of TRRAP and associated histone acetyl transferase complexes.[26, 33–35] At the core of MBII is a highly conserved four amino acid sequence (DCMW). W135 is highly conserved in MYC family proteins, [36, 37] and sits at the heart of the predicted MBII interface with TRRAP,[38] an interface that may be therapeutically targetable.[39]

MYC transforming activity is held in check by induction of apoptosis.[25, 40–43] Hence *MYC* deregulation in cancers is often accompanied by *TP53* inactivation or deregulated *BCL2*.[44–46] A range of aggressive B-cell neoplasms carry oncogenic events that arrest cells during differentiation between B-cell activation and PC differentiation.[47] MYC deregulation is a recurrent event in this context as a result of translocation or stabilizing mutations.[1, 3, 31, 32, 48, 49] Lymphomas with translocation of both *MYC* and *BCL2*, “double hit lymphoma”, as well as cases with MYC and BCL2 co-expression, without underlying translocation, “double expressing lymphoma”, have an adverse prognosis.[48, 50] Recently efficient transduction of primary human B-cells with oncogene combinations has provided a basis for *in vitro* modelling of human aggressive B-cell lymphoma.[51, 52] In this context MYC and BCL2 co-deregulation provided an example of a transforming combination driving sustained population expansion.[52]

Differentiation can oppose cellular transformation by driving cell cycle exit and limiting cellular plasticity. Concomitant with this MYC expression generally declines with differentiation.[15, 53, 54] We have developed models of human B-cell activation that are permissive for differentiation to a long-lived PC state.[55–57] Driven by signals mimicking antigen receptor ligation and T-cell help including transient CD40L exposure, B-cells undergo a process of cell growth and division in which endogenous MYC expression is first induced following activation and then repressed as the differentiating cells complete cell division and transition to a specialized secretory state, recapitulating physiological PC differentiation.[55] At the heart of the process of PC differentiation is a coordinated reorganization of transcription factors.[58] Overall the sequence of transcriptional regulation coordinates a MYC-associated burst of cell growth and division, with eventual repression of elements of the B-cell state and a switch to secretory gene expression.[7, 17, 58]

A trigger for the transition from growth program to PC differentiation is release from sustained CD40L signals,[59, 60] which provides potent NFκB pathway activation,[61, 62] and a signal which can delay and/or prevent differentiation of activated B-cells.[63–65] Sustained provision of CD40L was integral to previous *in vitro* modelling of human B-cell lymphomagenesis.[52] By sustaining CD40L signaling, the approach did not address whether oncogene deregulation sufficed to transform B-cells under conditions permissive for PC differentiation. Examining this is of interest because it would test the impact of deregulation of MYC in the context of an intrinsically reorganizing transcriptional program of differentiation. To address these questions, we have evaluated the impact of MYC deregulation in the context of BCL2 co-expression across human PC differentiation. Our data argue that MYC and BCL2 deregulation does not suffice to transform B-cells when conditions for differentiation are permissive, but that MYC diverts expression towards a distinct non-physiological pattern that alters metabolic and growth-related gene expression and impairs secretory output. These transcriptional impacts of MYC are independent of MBI but depend in part on MB0 and are dependent on MBII and the single amino acid W135.

## Results

### Acute MYC and BCL2 overexpression drives an aberrant B-cell differentiation phenotype

We aimed to test to what extent deregulated expression of MYC in combination with BCL2 impacted acutely on human B-cell differentiation. We initially evaluated a T58I variant of MYC in combination with BCL2, as this combination has been previously used in lymphoma modelling.[52] By including the T58I lymphoma-associated MYC stabilizing mutation in the MBI domain,[31, 32] this approach combined overexpression and stabilization to enhance the potential for MYC impact. We tested this in context of our differentiation system which is permissive for human B-cell differentiation to a long-lived PC state (Figure 1a). Briefly in this model system B-cells are activated by signals that include antigen receptor ligation, CD40 stimulation and cytokines IL2 and IL21 for 3 days, during which B-cells grow and begin to divide and endogenous MYC is expressed. At day 3 CD40 and antigen receptor ligation are removed and NFκB signalling is rapidly lost. Activated B-cells subsequently divide rapidly while transitioning to a plasmablast state. At day 6 plasmablasts are transferred to IL6 and APRIL or other cytokine containing conditions that support further differentiation to the PC state.[55–57] A significant difference from previous *in vitro* modelling of lymphomagenesis in which B-cells were continuously maintained in CD40 stimulating conditions,[52] is the removal of CD40 stimulation and NFκB activation at day 3 in our model supporting PC differentiation.[59] A distinct pattern of NFκB activation is subsequently reintroduced upon addition of APRIL at day 6 which supports differentiation to the PC stage.[57]

**Figure 1.**
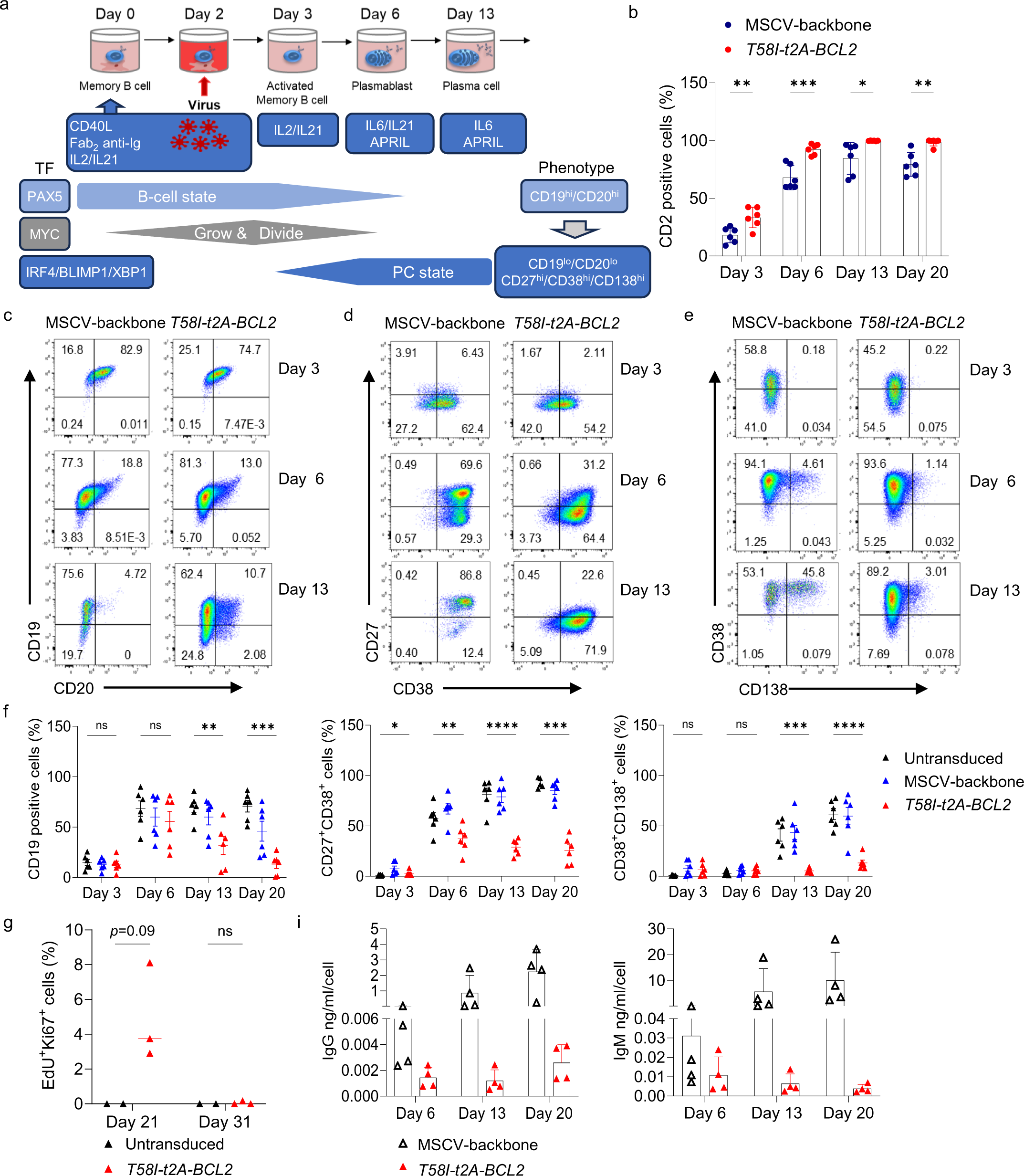
Acute MYC and BCL2 overexpression drives aberrant B cell differentiation. **a,** Graphical representation of the *in vitro* differentiation and transductions model system. This shows general phases by day of culture at the top, followed by summary culture conditions and biological processes below. Associated transcription factors (TF) are depicted on the left and phenotypic markers on the right. **b,** Flow cytometric quantitation of percentage CD2 positive cells for MSCV-backbone and *T58I-t2A-BCL2* conditions at indicated time points. **c, d, e,** Representative flow cytometry plots of CD19 vs CD20, CD27 vs CD38 and CD38 vs CD138, respectively, for MSCV-backbone and *T58I-t2A-BCL2* conditions at the indicated time points. **f,** Percentages of CD19 positive cells (left), CD27+CD38+ cells (middle), and CD38+CD138+ cells (right) for the indicated conditions at the indicated time points. **g,** Percentages of EdU+Ki67+ cells at the indicated time points for untransduced and *T58I-t2A-BCL2* samples after 1 hour of pulse EdU incorporation. Data shown for the MSCV-backbone and *T58I-t2A-BCL2* conditions are pre-gated to CD2+ populations (c, d, e, f, g) **i,** Quantification of total human IgG antibody secretion (left) and IgM antibody secretion (right) on day 6, day 13 and day 20 for the indicated conditions. Data are representative of at least two independent experiments. Bars and error represent mean and standard deviation (SD); Unpaired two-tailed Student’s *t-test*, (b, g). One-way ANOVA (f): ns, not significant; * *P* < 0.05; ** *P* < 0.01; *** *P* < 0.001; **** *P* < 0.0001.

In the context of this model peripheral blood memory B cells were transduced on day 2 of activation with *MYC T58I-t2A-BCL2* retroviral vector (henceforth *T58I-t2A-BCL2)* and were then returned to CD40 stimulating conditions for 24h before progressing into the differentiation protocol with removal of CD40 stimulation at day 3 (Figure 1a, Sup. Figure 1a). Transduction efficiency was high and sustained expression of the retroviral CD2 reporter was observed to day 20, by which time PC differentiation has been established for ten days in differentiations in the absence of transduction (Figure 1b).[56] Across multiple times points of differentiation, *T58I-t2A-BCL2* cells showed increased cell size relative to MSCV or untransduced controls (Sup. Figure 1c and d) accompanied by increased cell numbers at day 13 and day 20 (Sup. Figure 1b). *T58I-t2A-BCL2* was associated with a change in phenotype (Figure 1c-f). This included loss of CD27 and CD138 expression which are hallmark features linked to PC differentiation, while also showing a decrease in CD19 expression which is a hallmark B-lineage antigen whose loss of expression is a feature of malignant PCs. Thus, MYC T58I and BCL2 overexpression from the activated B-cell stage onward resulted upon subsequent differentiation in increased cell size and number and an aberrant phenotype with loss of typical PC markers.

### MYC and BCL2 overexpression delays cell cycle exit and secretory output

PC differentiation is characterized by cell cycle exit and reprogramming of the expression state toward secretory activity and away from cell growth and proliferation. MYC drives both cell growth and proliferation in lymphocytes.[5, 7, 17] We therefore assessed how MYC deregulation impacted on the proliferative status using a combination of EdU labelling and Ki67 detection at day 21 and 31 of differentiation (19 and 29 days, respectively, after transduction) in *T58I-t2A-BCL2* or control transduced cells. During differentiation of normal B-cells in the experimental culture system cell cycle exit occurs between day 6 and day 10 as the cells transition from plasmablast to PC state.[56] Consistent with the overall increase in cell number EdU^+^Ki67^+^ cells were readily detected in *T58I-t2A-BCL2* but not in control conditions at day 21. However, by day 31 *T58I-t2A-BCL2* conditions showed no significant increase in EdU^+^Ki67^+^ expressing cells (Figure 1g, Sup. Figure 1e). Therefore, under conditions permissive for B-cell differentiation to the PC state MYC and BCL2 deregulation resulted in an extended but ultimately curtailed proliferative state during subsequent differentiation.

During PC differentiation, proliferation and secretory reprogramming are coupled.[66–68] However, transition to a secretory state is not strictly linked to cell cycle exit. Both secreting and non-secreting cells have been reported to divide at similar rates.[69] We therefore tested secretory output of the *in vitro* differentiated cells. Secreted IgG and IgM normalized per cell, was significantly reduced in *T58I-t2A-BCL2* conditions at both day 6 and day 13 (Figure 1i). Thus, accompanying increased cell size and delayed cell cycle exit, MYC and BCL2 deregulation led to reduced secretory output from differentiating B-cells.

### Enforcing MYC overexpression drives classical target genes and alters patterns of transcription factor expression

To further understand the impact of MYC and BCL2 deregulation on differentiation we performed a time course gene expression study in control or MYC *T58I-t2A-BCL2* conditions. Gene expression was assessed at: day 0 - prior to activation; day 3 – activated B-cell stage, 24h after transduction; day 6 - plasmablast stage, 4 days after transduction; day 13 - early PC stage 11 days after transduction; and day 20 - established PC stage 18 days after transduction (MSCV-control conditions generated insufficient cells for analysis at day 20) (Sup. Table 1). Consistent with a progressive impact of MYC *T58I-t2A-BCL2* conditions on gene expression, uniform manifold approximation and projection (UMAP) showed similar clustering of samples at day 3 with subsequent increased separation of MYC *T58I-t2A-BCL2* conditions from controls at day 6, day 13 and day 20 (Figure 2a). MYC was substantially expressed in physiological differentiation at day 3 in activated B-cells and was then progressively repressed in control differentiation conditions. In contrast MYC *T58I-t2A-BCL2* conditions showed a modest increase in *MYC* expression at day 3 and then maintained supra-physiological levels of *MYC* expression throughout subsequent differentiation (Figure 2b). Expression of *CD2* and enhanced and sustained expression of *BCL2* was also confirmed in transduced conditions (Figure 2b and Sup. Figure 2a).

**Figure 2.**
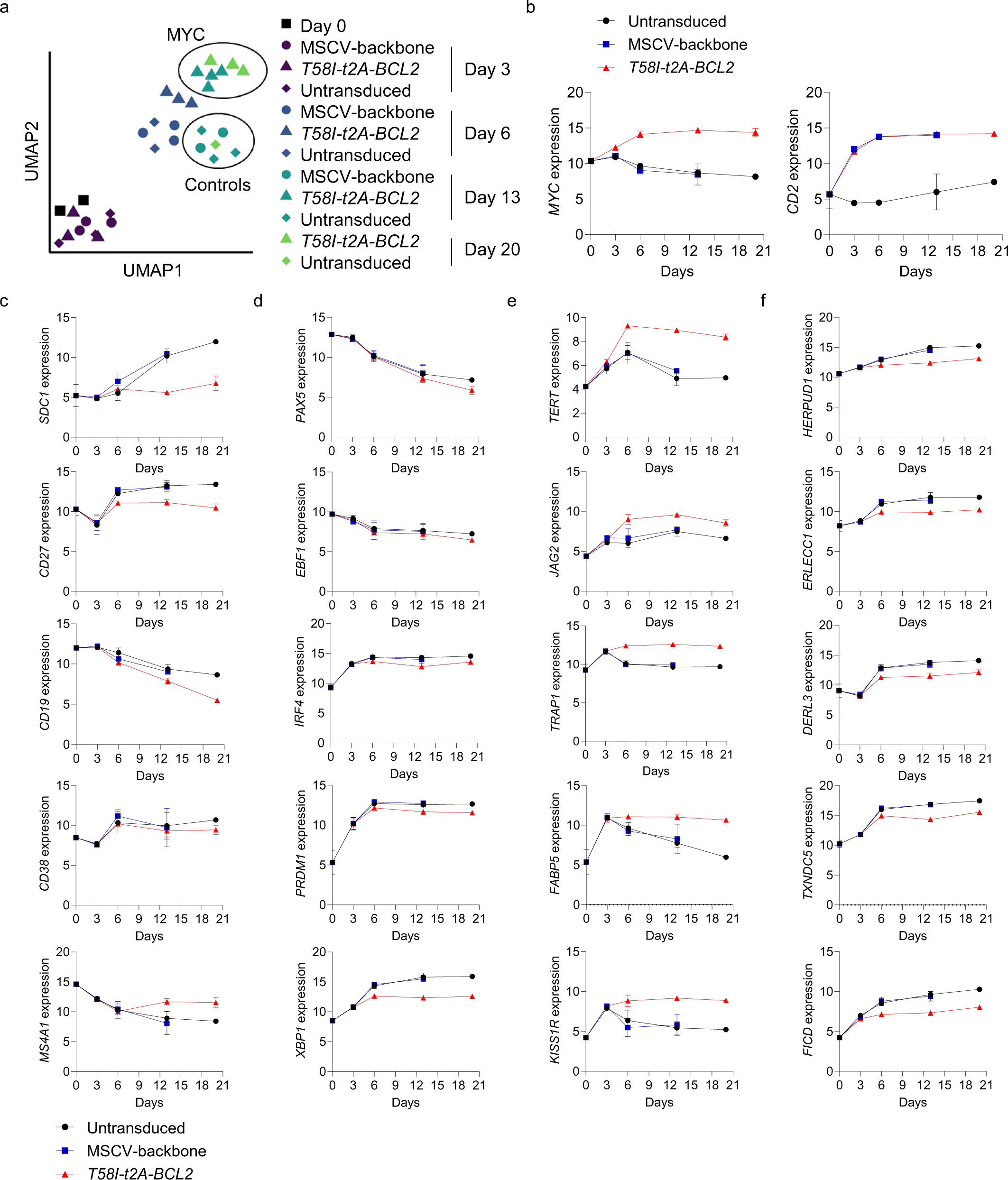
MYC T58I selectively perturbs gene expression in B-cell differentiation. **a,** Uniform Manifold Approximation Projection (UMAP) of differentially expressed genes for the indicated conditions and time points. **b, c, d, e, f,** Normalised RNAseq expression values (y-axis) of selected genes across the differentiation time course (x-axis days) as indicated for untransduced, MSCV-backbone and *T58I-t2A-BCL2* conditions. Gene expression is shown as indicated in the figure for **b**, *MYC* and *CD2*; **c**, surface proteins linked to immunophenotyping; **d**, transcription factors; **e**, known MYC targets; and **f**, XBP1 targets. Data are representative of two independent experiments with a total of n=1-4 samples per time point and condition.

We next assessed the extent to which phenotypic changes were recapitulated in gene expression data. We observed significant parallels with suppression of *SDC1* (CD138), *CD27* and *CD19* expression at later time points in MYC *T58I-t2A-BCL2* conditions relative to controls. *CD38* expression was not substantially impacted while *MS4A1* (CD20) expression was increased in MYC *T58I-t2A-BCL2* conditions (Figure 2c). These conditions were also associated with suppressed *TNFRSF17* (BCMA) but not *TNFRSF13B* (TACI) and *CD79A* but not *CD79B* expression (Sup. Figure 2b). Thus, the perturbed phenotype observed by flow cytometry was reflected in corresponding changes at transcript level.

PC differentiation is driven by coordinated changes in transcription factor expression.[58] We therefore examined how known regulators of the B-cell state and PC differentiation were expressed (Figure 2d). *PAX5* and *EBF1*, transcriptional regulators involved in maintaining B-cell states,[70–73] were expressed in resting and activated B-cells and then equivalently repressed during differentiation in control and MYC *T58I-t2A-BCL2* conditions. Positive regulators of PC differentiation *IRF4* and *PRDM1* (BLIMP1) were induced upon activation and differentiation with modest reductions observed in maximal expression for both factors in MYC *T58I-t2A-BCL2* conditions. *XBP1*, the primary transcriptional driver of secretory reprogramming and the unfolded protein response of the ER,[74–76] contrasted in showing more profoundly suppressed induction at day 6 and all subsequent time points in MYC *T58I-t2A-BCL2* conditions. Differential regulation of other transcription factors was also observed including repression of *RUNX1*, which has a role in cell cycle entry,[77] and enhanced expression of the PC fate antagonist *BACH2,*[78–80] and *SREBF1*, a controller of sterol metabolic pathways,[81] in MYC *T58I-t2A-BCL2* conditions (Sup. Figure 2c).

Consistent with canonical MYC-driven gene expression well-defined MYC targets such as *TERT*, the catalytic component of telomerase,[82] *JAG2*, a receptor on the NOTCH signalling pathway,[83] *TRAP1*, a key mitochondrial chaperone,[84] *FABP5*, linked to fatty acid metabolism and a potential therapeutic vulnerability in myeloma,[85] were increased (Figure 2e). A wide range of other MYC target genes defined in cellular models in both mouse and human[7] were also profoundly induced in MYC *T58I-t2A-BCL2* conditions.

Since XBP1 is a principal regulator of secretory reprogramming we assessed known XBP1 and ER stress response target genes. We found consistent patterns of repressed expression for genes such as *HERPUD1*, *ERLEC1, DERL3* and *TXNDC5*,[86–88] which share direct XBP1 promoter occupancy at the plasmablast stage in our model system (Figure 2f).[59] Immunoglobulin is the main secretory output of PCs and expression is profoundly decreased on conditional *XBP1* deletion in murine PCs.[89, 90] We therefore also examined the expression levels of immunoglobulin genes. Indeed, these genes comprised some of the most differentially expressed and showed significantly dampened expression particularly for *IGHG1*, *IGHG2*, *IGHG3* and *IGHM* in MYC *T58I-t2A-BCL2* conditions at later time points (Sup. Figure 2d). Thus, analysis at individual gene level suggested a coordinated impact of MYC overexpression on the regulation of metabolic and secretory pathways during PC differentiation and a separation of this perturbed secretory reprogramming from retained transcriptional control over features of the B-cell state.

### MYC and BCL2 overexpression drives coordinated modular patterns of gene expression change

To test gene regulation at a global level we analysed gene expression changes in MYC *T58I-t2A-BCL2* and control conditions using Parsimonious Gene Correlation Network Analysis (PGCNA).[91] This correlation-based method allows the shifting patterns of gene expression across differentiation and between conditions to be assessed in terms of modules of coregulated genes. PGCNA identified 16 modules of coregulated genes (labelled M1-M16 according to number of module genes (Sup. Table 2) with distinct patterns of expression across the differentiation and between control and MYC *T58I-t2A-BCL2* conditions (Figure 3a). Gene ontology and signature enrichment analysis demonstrated that the coregulated gene modules identified by PGCNA were highly significantly associated with different features related to B-cell differentiation, cell cycle, MYC function, translation, and metabolism (Figure 3b and Sup. Table 3). Module M1 was enriched for various features of the B-cell state including targets repressed by BLIMP and was expressed at day 0 and 3 and progressively repressed upon differentiation in a similar fashion in controls and *T58I-t2A-BCL2* expressing cells. Module M2 was enriched for genes regulated by NFκB reflecting the impact of the initial CD40:CD40L activation conditions, expression was low in day 0 B-cells, induced in day 3 activated B-cells and repressed on further differentiation in all conditions. In contrast to M1 and M2, the remaining 14 modules showed differential expression between controls and *T58I-t2A-BCL2* conditions. These separated between modules that: i) were repressed in normal differentiation but were enhanced in *T58I-t2A-BCL2* conditions (M4, M6 and M13), which were enriched for MYC regulated gene signatures; ii) modules that were induced upon differentiation but showed greater expression in control than *T58I-t2A-BCL2* conditions (M3, M5, M7, M16), including genes linked to XBP1 targets, UPR and ER secretory pathways; iii) modules that were induced upon normal differentiation but showed enhanced expression in *T58I-t2A-BCL2* conditions (M8, M10, M14), including genes linked to mitochondrial metabolism and OxPhos pathways; and iv) modules that were induced in control differentiation but repressed in *T58I-t2A-BCL2* conditions (M9, M12 and M15), including genes linked to autophagy and cellular quiescence.

**Figure 3.**
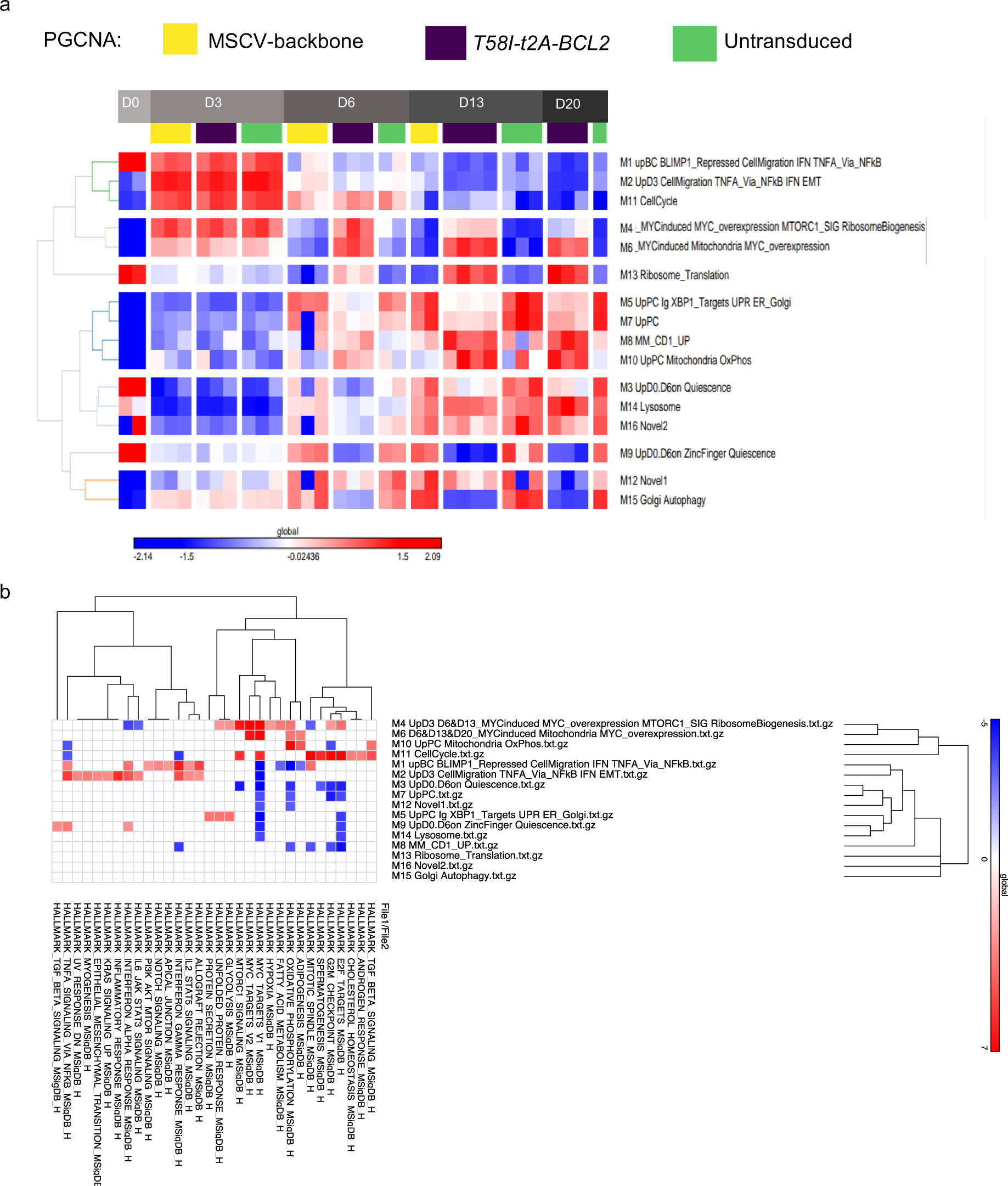
Expression network level analysis of MYC T58I on B-cell differentiation. Parsimonious Gene Correlation Analysis (PGCNA) was used to define modules of coregulated genes. **a,** Heat map of module level gene expression with expression patterns averaged across all genes per module on a z-score scale (−2.14 blue to +2.09 red). Modules are hierarchically clustered on the left and module number and indicative summary terms of associated ontologies are shown on the right. Time points are indicated in the grey to black bars at the top of the figure going from Day 0 (D0 left) to Day 20 (D20 right). Individual conditions are identified using the code as indicated at the top of the figure: yellow (MSCV-backbone), purple (*T58I-t2A-BCL2*), green (untransduced). **b,** Correlation plot of gene signature enrichment analysis for the 16 PGCNA-derived modules showing analysis for MsigDB Hallmark signatures. Enrichment is illustrated on a z-score scale from blue to red. With hierarchical clustering for enriched signatures and module enrichment patterns above and to the right. Module numbers and summary designations are identified to the right.

Modules linked to MYC regulated features (M4, M6 and M13) exhibited distinct expression patterns. Module M4 enriched for previously defined signatures of MYC overexpression, MTORC1 and ribosome biogenesis signatures were strongly induced by the activation conditions driving differentiation and were sustained in *T58I-t2A-BCL2* conditions at day 6 and to a lesser extent at day 13 but were largely repressed at day 20. Module M6 linked to MYC targets related to mitochondrial function was modestly induced by activation conditions at day 3 but was strongly induced in *T58I-t2A-BCL2* conditions at day 6 and sustained to day 20. Module M13 linked to a further group of genes related to ribosomes and translation, was modestly expressed in resting B-cells, repressed in all conditions upon activation and while further repressed upon differentiation in control conditions was super-induced beyond initial expression levels upon differentiation in *T58I-t2A-BCL2* conditions.

This time course analysis indicated that MYC T58I accompanied by BCL2 co-expression diverted differentiation toward a distinct expression state, driving modules of metabolic and translation related genes that were not physiologically expressed in PC differentiation. At the same time, while suppressing features of the PC secretory state, MYC T58I did not interfere with repression of the B cell-state during plasma cell differentiation. The impact of overexpressed MYC was not fixed across the time course, nor did it reflect an amplification of the physiological differentiation programs. Instead, the impact of MYC developed across the course of the differentiation toward a distinct aberrant expression state.

### MB domains of MYC show differential contributions to gene regulation during PC differentiation

The MYC TAD has been implicated as critical in transforming activity in various cellular models, we therefore next aimed to test the contribution that MB0, MBI or MBII domains of the MYC TAD made to the divergent programming of perturbed PC differentiation. To this end we generated a MYC *WT-t2A-BCL2* vector (i.e. with T58 not T58I) and vectors in which MB0, MBI or MBII were deleted in this context (Sup. Figure 3a). We obtained similar transduction efficiencies to our previous results (Sup. Figure 3b). MYC *WT-t2A-BCL2* recapitulated the phenotypic effect observed for MYC T58I. *ΔMBI-t2A-BCL2* which removes the phosphodegron sequence encompassing T58 showed little difference in phenotype from MYC *WT-t2A-BCL2*. By contrast *ΔMB0-t2A-BCL2* showed partial reversal and *ΔMBII-t2A-BCL2* showed significant reversal of the MYC-associated phenotype and reversion toward phenotypic patterns of control conditions (Figure 4 a-c, Sup. Figure 3e). Overexpression of MYC and BCL2 proteins in comparison to the MSCV-backbone control was validated for all three MYC MB deletion mutants tested, including the enhancement of MYC expression expected for *ΔMBI-t2A-BCL2* with phosphodegron sequence removal. Similar levels of BLIMP1 expression were observed following differentiation in all conditions (Sup. Figure 3c, d).

**Figure 4.**
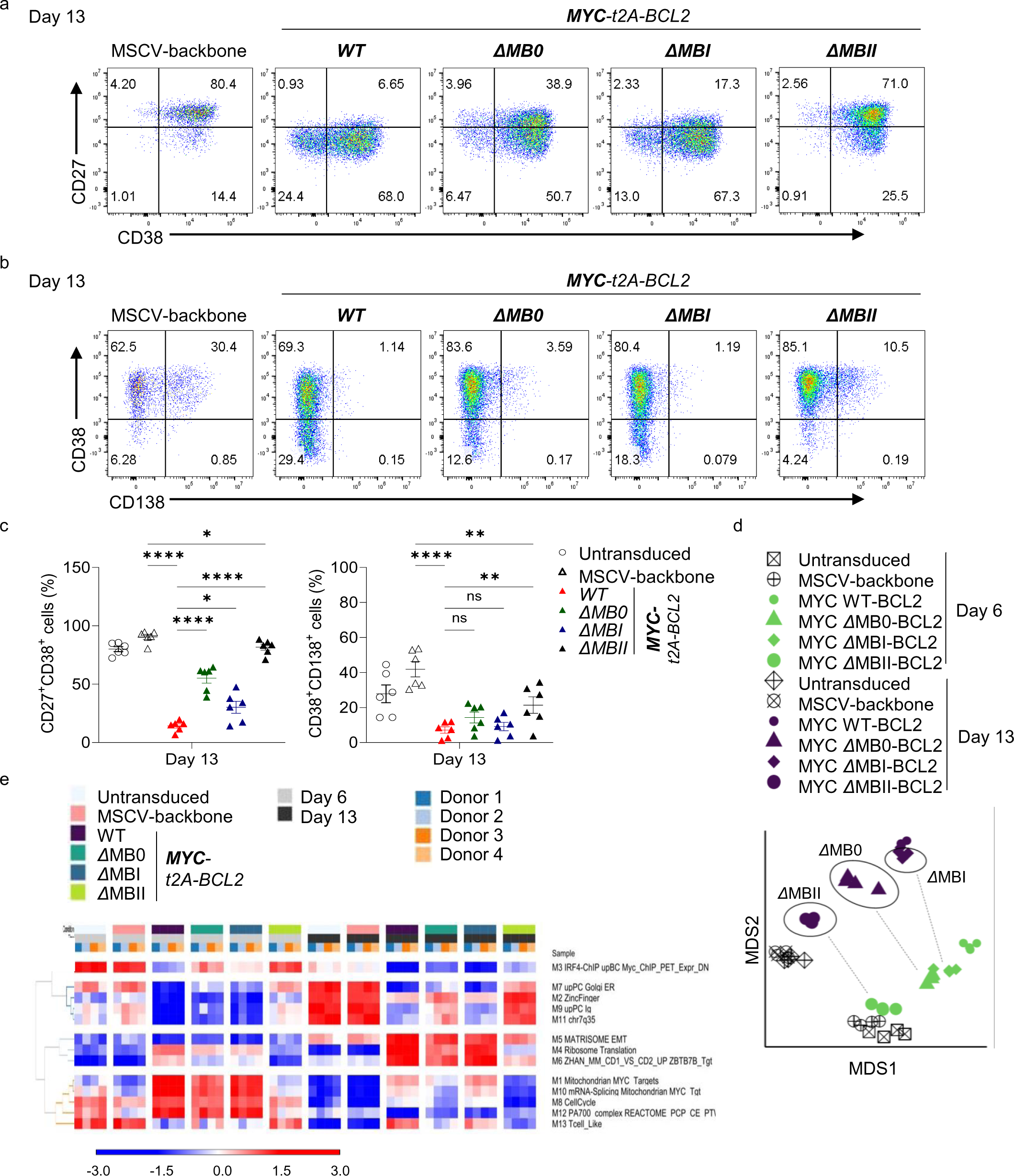
Deletion of MYC TAD MB0, MBI and MBII have differential effects on MYC driven phenotypic and expression features. **a, b,** Representative flow cytometry plots at day 13 for control MSCV-backbone, MYCwt, ΔMB0, ΔMBI and ΔMBII conditions as shown for **a**, CD27 vs CD38, and **b**, CD38 vs CD138. **c,** Summary of flow cytometrically defined percentages of CD27+CD38+ cells (left) and CD38+CD138+ cells (right), at day 13 for the indicated conditions. Data are representative of three independent experiments. Bars and error represent mean and standard deviation (SD); Unpaired two-tailed Student’s *t-test*: ns, not significant; * *P* < 0.05; ** *P* < 0.01; **** *P* < 0.0001. Data shown for the transduced conditions are pre-gated to CD2+ populations (a, b, c,). **d,** Multidimensional Scaling (MDS) of differentially expressed genes at the day 6 (green) and day 13 (purple) time points, and controls, for the indicated samples as illustrated in the figure. **e,** PGCNA defined modules of coregulated genes shown as a heat map of module level gene expression with expression patterns averaged across all genes per module on a z-score scale (−3 blue to +3 red). Modules are hierarchically clustered on the left and module number and indicative summary terms of associated ontologies are shown on the right. Time points are indicated as grey day 6 and black day 13. Samples from different donors are illustrated in the blue to orange color code, and for individual conditions with color code identified in the figure.

To further assess the impact of MB deletions, we studied gene expression at day 6 (plasmablast) and day 13 (PC stage) when divergent patterns of expression changes were observed in MYC *T58I-t2A-BCL2* expressing cells (Sup. Table 4). Multidimensional scaling (MDS) showed that the distinct conditions separated in terms of differentiation – separating control differentiations at day 6 from those at day 13 – and according to MYC transduction at each time point (Figure 4d). This was consistent with the hierarchal order of phenotypic impact such that MYC *WT-t2A-BCL2* was most distinct from controls at both day 6 and day 13. *Δ*MBI shifted from being most similar to *Δ*MB0 at day 6 to become most similar to MYCwt at day 13. *Δ*MB0 fell between MYCwt and control differentiation conditions at both day 6 and day 13, and *Δ*MBII clustered in closest proximity to the controls at both time points but was more divergent at day 13 than day 6 (Figure 4d). We next assessed changes at modular level. PGCNA again successfully resolved modules of genes enriched for biological processes related to MYC function (M1, M4, M8 and M10) and PC differentiation (M7 and M9) that were differentially expressed between time points (Figure 4e and Sup. Table 5). MYCwt and *Δ*MBI showed similar patterns of module regulation that diverged from those observed in controls, with enriched biological processes mapping onto those observed for differential regulation by MYC T58I conditions (Sup. Table 6). *Δ*MB0, while similar to MYCwt and *Δ*MBI, diverged in the level of intensity of expression of both MYC up- and down-regulated modules. In contrast, *Δ*MBII showed markedly reduced evidence of MYC-associated module regulation and showed greater expression of PC-associated modules than other MYC conditions (Figure 4e). Therefore, both in the context of dimensionality reduction using MDS and modular expression analysis using PGCNA, the MB deletion mutants showed consistent features with a hierarchy of impact on the consequences of MYC overexpression: *Δ*MBII > *Δ*MB0 > *Δ*MBI.

To consider this at a gene level we revisited the expression patterns of index genes linked to B-cell differentiation, MYC impact and XBP1 function. Here we observed that while MYC and CD2 expression levels were similar in all conditions (Sup. Figure 4), a hierarchy was evident across both MYC responsive and repressed features at individual gene level for surface proteins, transcription factors, MYC targets, UPR/ER genes and immunoglobulin genes (Figure 5a-d and Sup. Figure 4).

**Figure 5.**
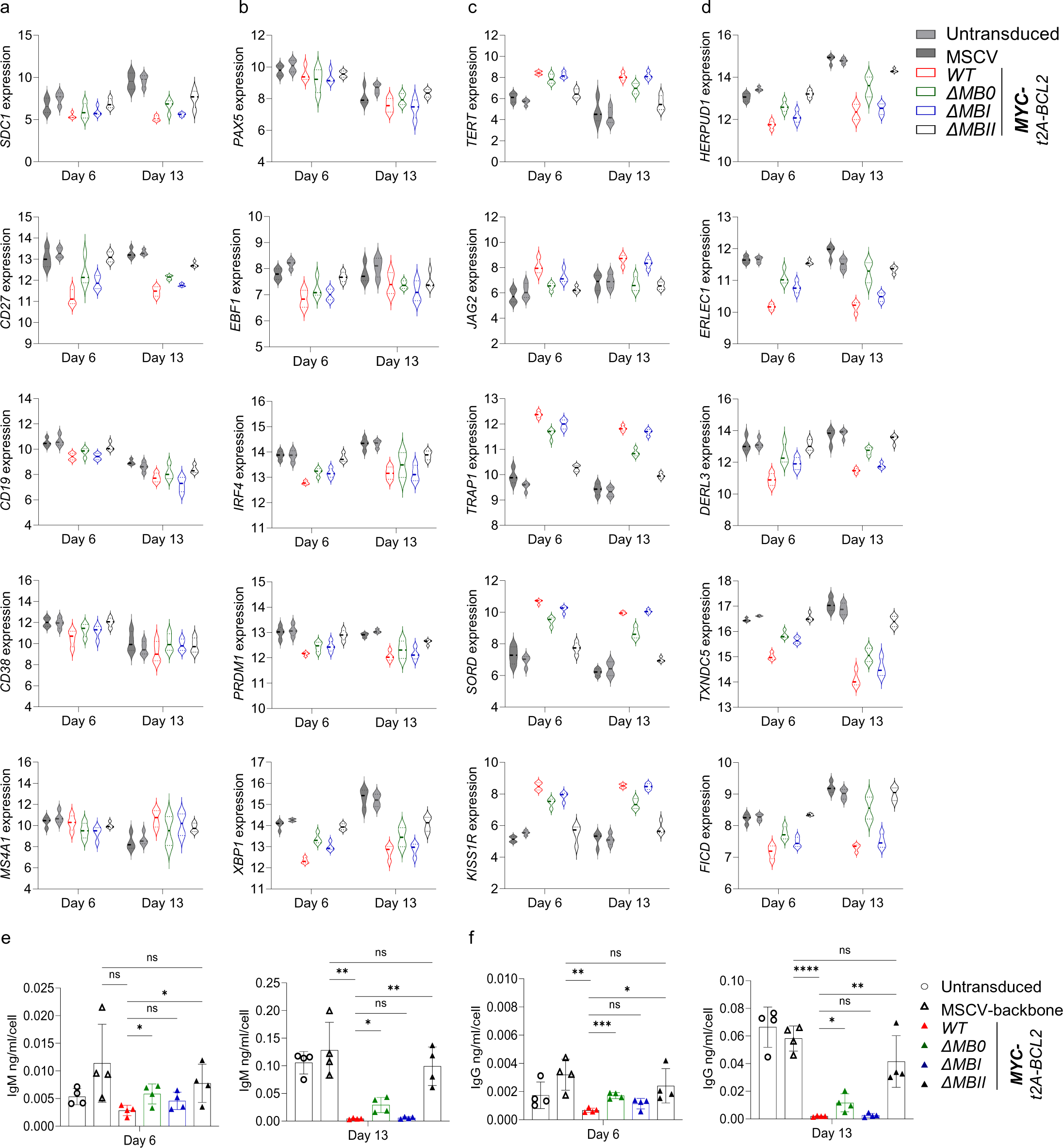
MB0, MBI and MBII deletion impacts on indicative gene regulation and on functional secretory output. Violin plots of log2 normalised RNAseq expression values of individual genes plotted at day 6 (left side of graphs) and day 13 time points (right side of graphs) for the indicated conditions (top right of figure). Genes shown are indicated to the left of each graph for: **a**, surface antigens; **b**, transcription factors; **c**, MYC targets; and **d**, XBP1 targets. Data are representative of two independent experiments with a total of n=4 samples per time point and condition. Quantification of **e**, IgM and **f**, IgG antibody concentration normalized per cell at day 6 (left graph) and day 13 (right graph) for conditions as indicated to the lower right of the figure. Data are representative of two independent experiments. Bars and error represent mean and standard deviation (SD); Unpaired two-tailed Student’s *t-test*: ns, not significant; * *P* < 0.05; ** *P* < 0.01; *** *P* < 0.001; **** *P* < 0.0001.

The observed changes in secretory and immunoglobulin gene expression suggested that functional secretory activity would be differentially impacted. Supernatants were tested by ELISA for secreted IgM and IgG at day 6 (Figure 5e) and day 13 (Figure 5f) from the distinct conditions. Concordant with gene expression, secretory output was detected at day 6 and rose around 10-fold by day 13 in control conditions. Secretory rates were similarly repressed in MYCwt and *Δ*MBI conditions, with repression most evident at the later time point when high output is established under control conditions. Secretory output remained substantially reduced in *Δ*MB0 conditions but significantly higher than MYCwt, while for *Δ*MBII conditions the MYC effect was lost and there was no significant difference from control conditions.

### The DCMW motif and W135 are critical for the effect of MYC MBII on human PC differentiation

Given the apparent dependence on MBII we next addressed to what extent this could be attributed to the core conserved sequence of MBII: the 132-135 aa DCMW motif or W135 alone, the most highly conserved residue in MBII which sits at the heart of the predicted TRRAP interaction.[38] Substitutions DCMW/AAAA or W135A were generated in the context of the *MYC-t2A-BCL2* configuration (Sup. Figure 5a) and comparably high transduction efficiency was verified for the MBII mutants (Sup. Figure 5b). MYC, BCL2 and BLIMP1 protein overexpression was validated (Sup. Figure 5c, d). BCL2 and BLIMP1 expression was equivalent between conditions. MYC was substantially overexpressed relative to controls in all conditions but was significantly higher in MYCwt relative to *Δ*MBII or MBII point mutants suggesting that MBII may in part contribute to stability in this context (Sup. Figure 5d). The MYC *MBII-4aa* mutant or the *MBII-W135A* again significantly impacted on the phenotypic changes induced by MYC, such that MYC *MBII-4aa* or *MBII-W135A* expressing cells more closely resembled control differentiations than MYCwt conditions (Figure 6a-c and Sup. Figure 5e). Our previous analyses indicated that phenotypic effects of MYC conditions were closely related to the extent of gene expression change in the model. Indeed, in multidimensionality scaling of day 13 gene expression data *Δ*MBII and the two MBII mutants clustered together with untransduced controls and separate from MYCwt conditions (Fig. 6d). Absolute numbers of significantly differentially expressed genes even at lenient fold-change thresholds were profoundly reduced for *Δ*MBII and the two MBII mutant conditions (Sup. Table 7). PGCNA confirmed that at modular level MYCwt conditions again differed profoundly from controls (Fig. 6e) with consistent biological differences to previous results (Sup. Table 8 and 9). While in *Δ*MBII and the MBII mutant conditions expression of modules related to PC differentiation and immunoglobulin gene expression were restored (Fig. 6e), modules linked with MYC-associated genes were less induced (Fig. 6e). A similar pattern was observed for genes indicative of MYC impact (Supplemental Figure 6). Indeed no significantly differentially expressed genes were observed between *Δ*MBII and either MBII mutant or between the MBII mutant conditions in direct comparison. When comparing *Δ*MBII or either of the two MBII point mutant conditions to untransduced control conditions only small numbers of genes remained differentially expressed. *XBP1* remained modestly reduced relative to control conditions but was also significantly restored relative to MYCwt (Figure 7a). Potential XBP1 target genes including immunoglobulins genes were not significantly different between *Δ*MBII or MBII mutants and control conditions indicating that function of secretory reprogramming was retained under these conditions (Figure 7a). Similarly, almost all MYC induced gene expression changes were lost in the context of *Δ*MBII or MBII substitutions with only very few genes showing retained significant expression differences (Figure 7b and Sup. Figure 6a). Consistent with the similarity between *Δ*MBII and MBII mutant conditions at gene expression level, ELISA demonstrated comparable IgM and IgG secretion in these conditions relative to control at day 6 and day 13 while in MYCwt conditions the estimated per cell secretory output was again profoundly repressed at day 13 (Figure 7c and d). We conclude that either a four amino acid DCMW/AAAA substitution or a single W135A substitution sufficed to phenocopy deletion of MBII and abrogate features associated with MYC overexpression in human B-cell differentiation.

**Figure 6.**
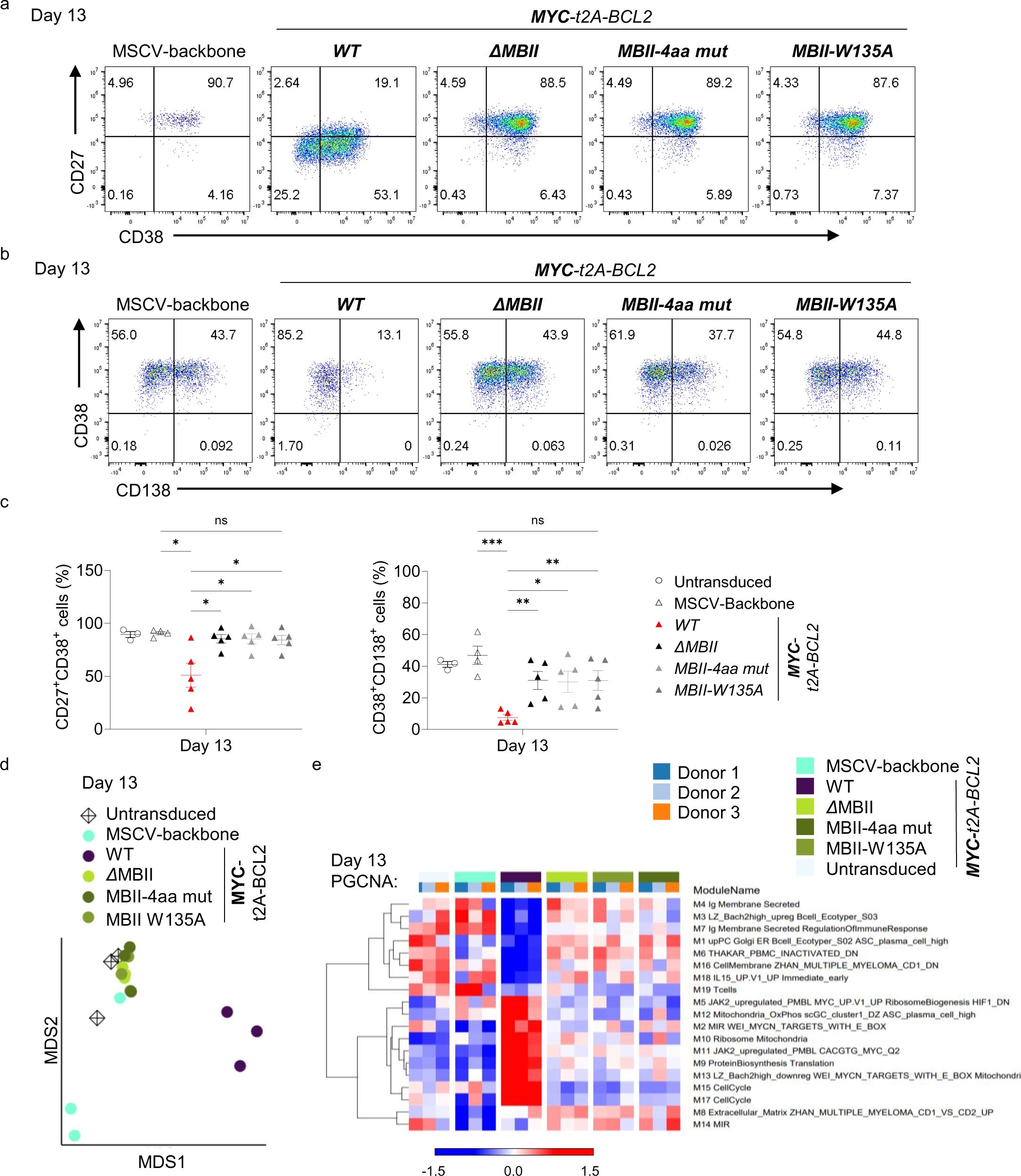
Point mutation of the DCMW motif and W135 phenocopy MBII deletion. Representative flow cytometry plots at day 13 for the indicated conditions above each dot plot for: **a,** CD27 vs CD38; and **b,** CD38 vs CD138. **c,** Flow cytometric quantification of percentage data CD27+CD38+ cells (left), and CD38+CD138+ cells (right) at day 13 for the conditions indicated to the right of graphs. Data are representative of at least three independent experiments. Bars and error represent mean and standard deviation (SD); Unpaired two-tailed Student’s *t-test*: ns, not significant; * *P* < 0.05; ** *P* < 0.01; *** *P* < 0.001. Data shown for the conditions tested, apart from the untransduced, are pre-gated to CD2+ populations (a, b, c). **d,** Multidimensional Scaling (MDS) of differentially expressed genes at day 13 for the indicated samples. **e,** PGCNA defined modules of differentially coregulated genes at day 13 shown as a heat map of module level gene expression with expression patterns averaged across all genes per module on a z-score scale (−1.5 blue to +1.5 red). Modules are hierarchically clustered on the left and module number and indicative summary terms of associated ontologies are shown on the right. Samples from different donors are illustrated in the blue to orange color code and for individual conditions with color code identified above the figure.

**Figure 7.**
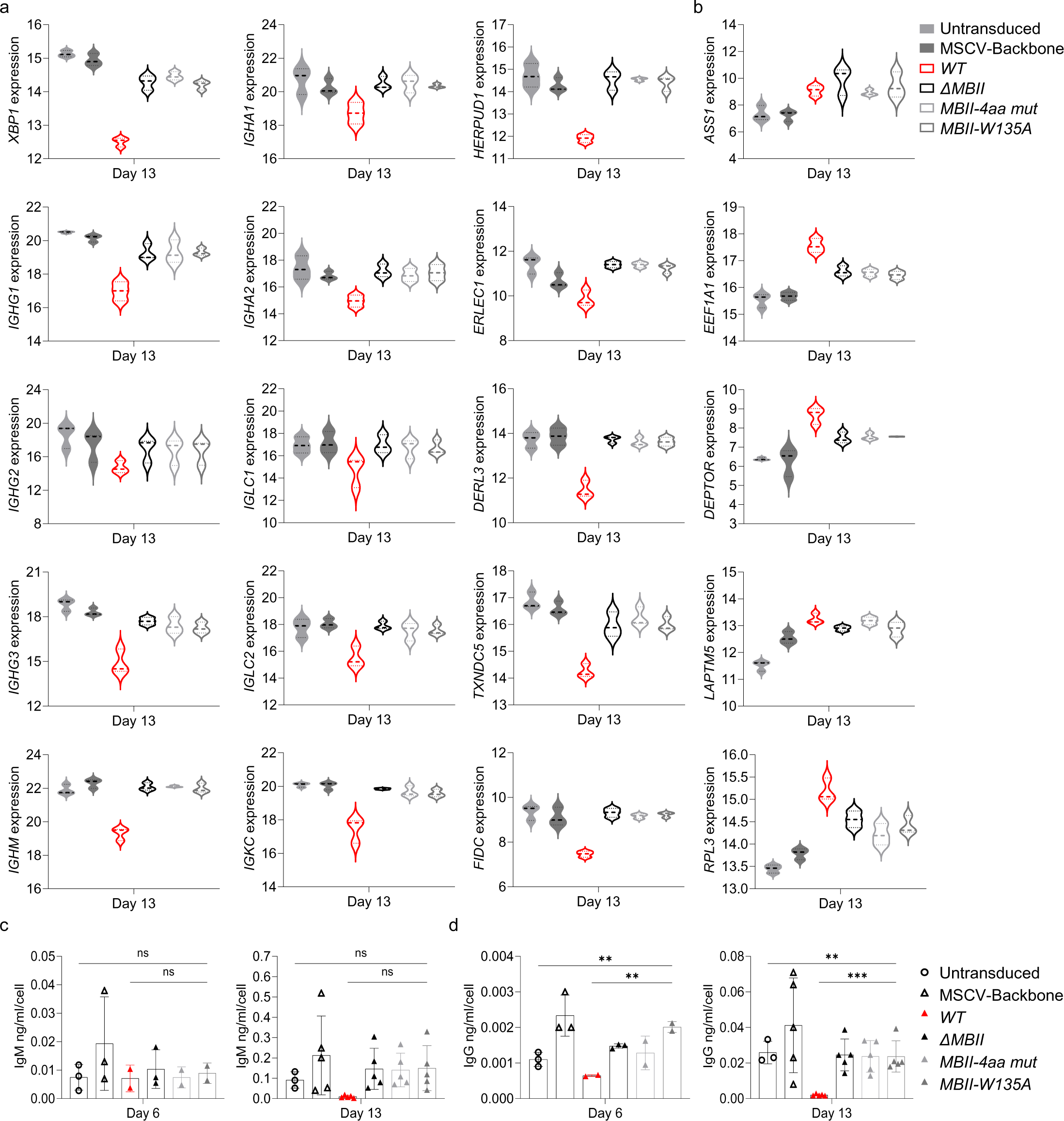
Point mutation of the DCMW motif and W135 phenocopy MBII deletion and retain only minimal impact on secretory reprogramming and MYC target gene regulation. Violin plots of log2 normalised RNAseq expression values of genes identified on the y-axis of each graph at day 13 for the conditions as indicated to the top right of the figure: **a,** XBP1, immunoglobulin genes and XBP1 targets; and **b,** MYC targets. Data are representative of two independent experiments with a total of n=3 samples per time point and condition. Quantification of **c,** IgM and **d,** IgG, antibody concentration normalized per cell at day 6 (left graph) and day 13 (right graph). Data are representative of two independent experiments. Bars and error represent mean and standard deviation (SD); One-way ANOVA: ns, not significant; ** *P* < 0.01; *** *P* < 0.001.

## Discussion

Deregulation of MYC is one of the key drivers of aggressive B-cell malignancies.[3, 4, 92] At the same time precise control of MYC expression is essential to co-ordinate cell growth and proliferation with differentiation in functional lymphocyte expansion of the immune response.[5–7] Here we have addressed the acute impact of MYC overexpression during human B-cell differentiation to the PC stage. We have assessed the extent to which MYC overexpression prevents or perturbs PC differentiation and the extent to which the impact of MYC overexpression changes as the cellular context progresses along the differentiation trajectory. We tested this both for MYC carrying the T58I mutation found in aggressive lymphoma, which stabilizes MYC expression, and wild type MYC lacking this mutation.[29, 32, 93] MYC deregulation drove gene expression changes consistent with previous models of lymphoma and other cancer types.[7] However the effect of MYC overexpression changed as the underlying cellular differentiation state shifted towards the PC state.

Distinct models of gene regulation by overexpressed MYC have been proposed, suggesting either action as a global transcriptional activator or as a more specific regulator of distinct biological pathways.[11, 15–17] In our model MYC expression is sustained at high levels from the activated B-cell stage onward and correlates with progressively lower endogenous MYC levels in control conditions. Our experiments included BCL2 co-expression to provide rescue from potential MYC driven apoptosis,[45, 46] and did not directly distinguish the potential effects of BCL2 over-expression alone. However, the various MYC mutants demonstrated that the gene expression changes depended on elements of MYC and the impact of BCL2 on gene expression was negligible. We can therefore assign the distinct expression states to MYC activity with reasonable confidence. Genes that were enhanced or suppressed by overexpressed MYC during differentiation were significantly linked to known biology and enriched for hallmark MYC targets genes and those with classical E-box motifs. However, the expression patterns linked to MYC overexpression were neither constant over the course of differentiation nor readily explained as enhanced expression of the prevailing patterns of gene expression in the control differentiations. Rather overexpressed MYC established a distinct expression state during PC differentiation.

Physiologically MYC has been identified as a rheostat linking the extent of B-cell activation to cell growth and the subsequent capacity for sequential cell division.[6–8] In this setting MYC sits at the heart of a transcriptional circuitry which coordinates a burst of cell growth and division with eventual differentiation and secretory reprogramming (Supplemental Figure 7a).[58] During this sequence of events external signals drive B-cell activation and expression of MYC.[7, 94–96] These signals also induce expression of IRF4 which in turn cooperates with STAT3 to drive expression of PRDM1/BLIMP1.[97, 98] Accumulating expression of BLIMP1 ultimately leads to repression of *MYC*,[20] curtailing the proliferative burst. At the same time BLIMP1 suppresses the B-cell identity transcription factor *PAX5* and other features linked to the mature B-cell state.[70, 71] Loss of PAX5 releases *XBP1* from PAX5 mediated repression.[23, 71, 99] Acting together BLIMP1 and XBP1 co-ordinate the high-level expression of immunoglobulin genes, the transition from membrane to secreted immunoglobulin encoding transcripts, and the expansion of the secretory apparatus.[23, 74–76, 89, 90, 100] The suppression of *MYC* and expression of negative cell cycle regulators coordinates the secretory transition to cell cycle exit.[21, 101, 102] Additional transcription factors such as BACH2 and BCL6 can add further levels of control to delay differentiation and facilitate class switching or interpose the complex biology of the germinal centre response.[78–80, 103–106]

The enforced expression of MYC that we have modelled occurs at the critical juncture when the burst of physiological MYC expression is at its peak and then begins to be curtailed by the underlying reorganizing transcription factor network.[59] Here deregulated MYC expression sustains gene expression linked to the cell growth and metabolic programs. Importantly deregulated MYC expression has little impact on the expression of *IRF4* or *BLIMP1* or the transcriptional regulators of B-cell identity *PAX5* or *EBF1*. Hence the regulatory circuitry controlling changes in B-cell identity related genes is at most marginally affected.[22, 70–73] This contrasts with a profound impact of MYC deregulation on the initiation of secretory output and secretory reprogramming, which is coupled with dampening of *XBP1* expression and suppression of immunoglobulin gene enhancement. MYC has previously been identified as capable of binding to the *XBP1* promoter and positively regulating expression in cancer models.[107, 108] Indeed, MYC is often deregulated in aggressive PC malignancies in which XBP1 is expressed.[109] The contrast with the suppressive effect of MYC overexpression on *XBP1* and secretory programming that we have observed in B-cell differentiation could be explained by differential recruitment of cofactors by MYC in this setting. Such differential recruitment would have parallels in the complex web of potential MYC interactions observed across other cell systems.[18] We have not addressed to what extent the delay in secretory programming is driven by repression of *XBP1* as opposed to an impact on immunoglobulin gene expression and protein load. However, murine conditional knockout models argue for a critical role of XBP1 in transcriptional enhancement of immunoglobulin gene expression.[90] Hence, we favor a model in which deregulated MYC suppresses *XBP1* which in turn leads to reduced immunoglobulin gene expression and delay in secretory reprogramming. This regulatory arrangement would be consistent with the role of MYC in the phase of lymphocyte activation and growth prior to differentiation,[5–8, 17] with MYC deregulation simultaneously biasing gene expression toward anabolic pathways and away from pathways associated with export of cellular material as secreted immunoglobulin. If the transcriptional circuitry controlling PC differentiation is viewed as two interconnected feedforward loops, one regulating the pulse of growth and proliferation and the other the delayed transition to a secretory fate, then the deregulated overexpression of MYC in activated B-cells disrupts differentiation by uncoupling both of these interconnected feedforward loops (Supplemental Figure 7b).

The MYC TAD harbors evolutionarily conserved amino acid sequences essential for MYC-mediated transformation.[18, 26, 27] Our data demonstrate a similar dependence on MB domains for the transcriptional impact of MYC during B-cell differentiation. In this setting *Δ*MBI resulted in relatively enhanced MYC protein expression similar to T58I and consistent with ablation of the phosphodegron sequence.[30] *Δ*MB0 led to a partial suppression of *XBP1* and secretory reprogramming and a reduced ability to induce a select subset of genes positively regulated by MYCwt. By contrast *Δ*MBII or selective amino acid mutations, including the single point substitution W135A, rendered MYC all but nonfunctional. The MBII domain is particularly notable for recruitment of TRRAP and associated histone acetyl transferase complexes.[26, 33, 34] The DCMW motif and W135 sit at the heart of the predicted interface with TRRAP.[38] Nevertheless, the degree to which MBII deletion, mutation of DCMW, or W135 impacted was unexpected. While the MBII domain is critical for MYC mediated transformation in fibroblasts, the W135A mutation had little effect in this context and an W135E substitution was needed to phenocopy MBII deletion.[35] MBII mutants can bind to physiological MYC targets in U2OS cells.[37] Furthermore, *Δ*MBII MYC could partially compensate for MYCwt in drosophila development.[110] That all three versions of MYC targeting MBII in our model produce the same effect provides evidence for a critical dependence on MBII in the context of MYC deregulation during PC differentiation. Other studies have identified the interaction of MYC with WDR5 via the MBIIIb region of the MYC TAD as critical for recruitment of MYC to chromatin, including in Burkitt lymphoma.[111–114] The MBIIIb region along with the MYC DNA binding domain are intact in *Δ*MBII and MBII mutants. However, it remains to be tested whether the profound impact on overexpressed MYC in B-cell differentiation is explained by a requirement for MBII to support recruitment of excess MYC to target genes or to drive subsequent gene regulation. A recent study has shown that in U2OS cells high level MYC stabilizes and extends long-range chromatin interactions.[115] If MYC DNA binding is retained in the MBII mutants, a possible explanation for the global impact of MBII mutants could be an essential role for MBII in mediating such chromatin effects.

In summary we have tested a model that allows the acute effect of MYC overexpression in cooperation with BCL2 to be studied as human B-cells differentiate to the PC state. This demonstrated that MYC overexpression did not transform B-cells under conditions permissive for differentiation. Instead, MYC overexpression selectively impacted on the metabolic and secretory components of reprogramming, leaving the element of differentiation associated with repression of the B-cell state largely intact. The TAD domain MB0 and MBII were necessary for MYC effects, and a critical dependence on MBII could be resolved to the DCMW motif and W135.

## Data Availability Statement

The primary datasets are available at the Gene Expression Omnibus GSE262809

## Acknowledgements

We thank Erica Wilson for technical support and advice and Ulf Klein and Richard Bayliss for advice, support and critical review of this work.

## Funding Acknowledgements

This work was supported by the Ella Dickinson Charitable Foundation Scholarship Program (scholarship recipients: P.V., B.K., A.M.; investigators: R.O., G. D., R.T.) and an Intercalated Degree 1022 Award, Pathological Society of Great Britain and Northern Ireland and EXSEL scholarship, University of Leeds School of Medicine to E.P.. This work was supported by Cancer Research UK program grant (C7845/A29212) (M.C, G.D., and R.T) and Cancer Research UK and FC AECC and AIRC under the Accelerator Award Program (C355/A26819). This research is funded in part (M.C.) by the National Institute for Health and Care Research (NIHR) Leeds Biomedical Research Centre (BRC) (NIHR203331). R.T., G.D., are supported in part by the National Institute for Health and Care Research (NIHR) Leeds Biomedical Research Centre (BRC) (NIHR203331). D.J.H. was supported by a fellowship from Cancer Research UK (CRUK) (RCCFEL∖100072) and received core funding from Wellcome (203151/Z/16/Z) to the Wellcome-MRC Cambridge Stem Cell Institute and from the CRUK Cambridge Centre (A25117). D.J.H is supported by the National Institute for Health and Care Research (NIHR) Cambridge Biomedical Research Centre (BRC-1215-20014). The views expressed are those of the authors and not necessarily those of the NIHR or the Department of Health and Social Care. For the purpose of Open Access, the authors have applied a CC BY public copyright licence to any Author Accepted Manuscript version arising from this submission.

