## Supplementary Figure 1 for "Enforced MYC expression selectively redirects transcriptional programs during human plasma cell differentiation"

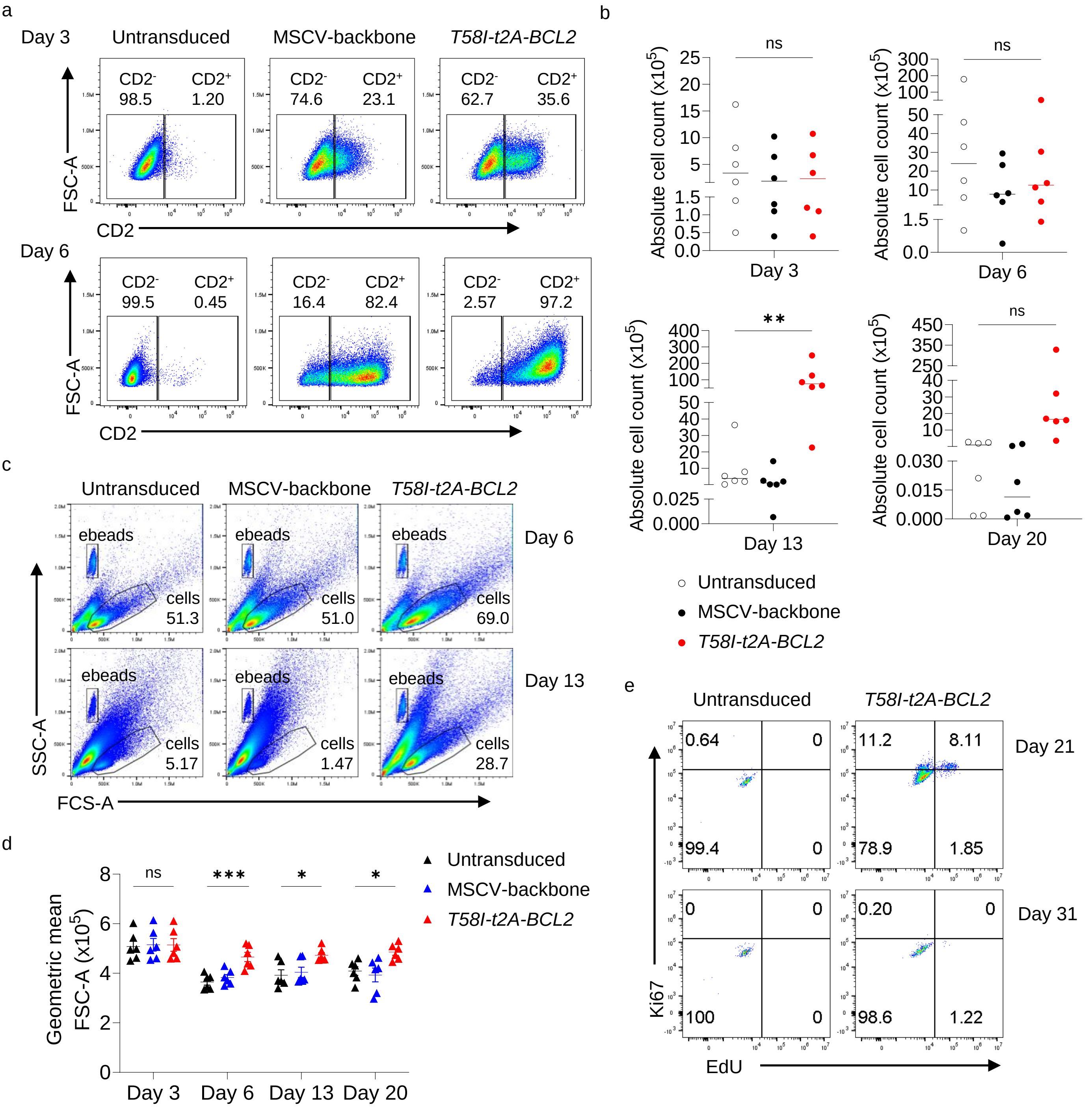


**Supplementary Figure 1. MYC-BCL2 overexpression results in increased cell number and size and extends the proliferative window. (Accompanies Figure 1) a,** Representative flow cytometry plots of forward scatter on area parameter (FSC-A) against CD2 showing transduction efficiency at days 3 and 6 for the indicated conditions. **b,** Cell counts at day 3, 6, 13, and 20 for the indicated conditions. **c,** Flow cytometry data illustrating side versus forward scatter parameters on area, SSC-A and FSC-A at day 6 (top) and day 13 (bottom) for the indicated conditions. Counting beads labeled as ebeads along with viable cell gating. **d,** Calculated geometric mean of FSC-A data for the indicated time points and conditions. Bars and error represent mean and standard deviation (SD); One-way ANOVA (b,d): ns, not significant; * $P < 0.05$; ** $P < 0.01$; *** $P < 0.001$. **e,** Representative flow cytometry plots of Ki67 versus EdU after 1 hour pulse incorporation conducted at day 21 and day 31 of the conditions indicated above the figure. *T58I-t2A-BCL2* condition is pre-gated to the $CD2^+$ population. Data are representative of three independent experiments.
