## Supplementary Figure 2 for "Enforced MYC expression selectively redirects transcriptional programs during human plasma cell differentiation"

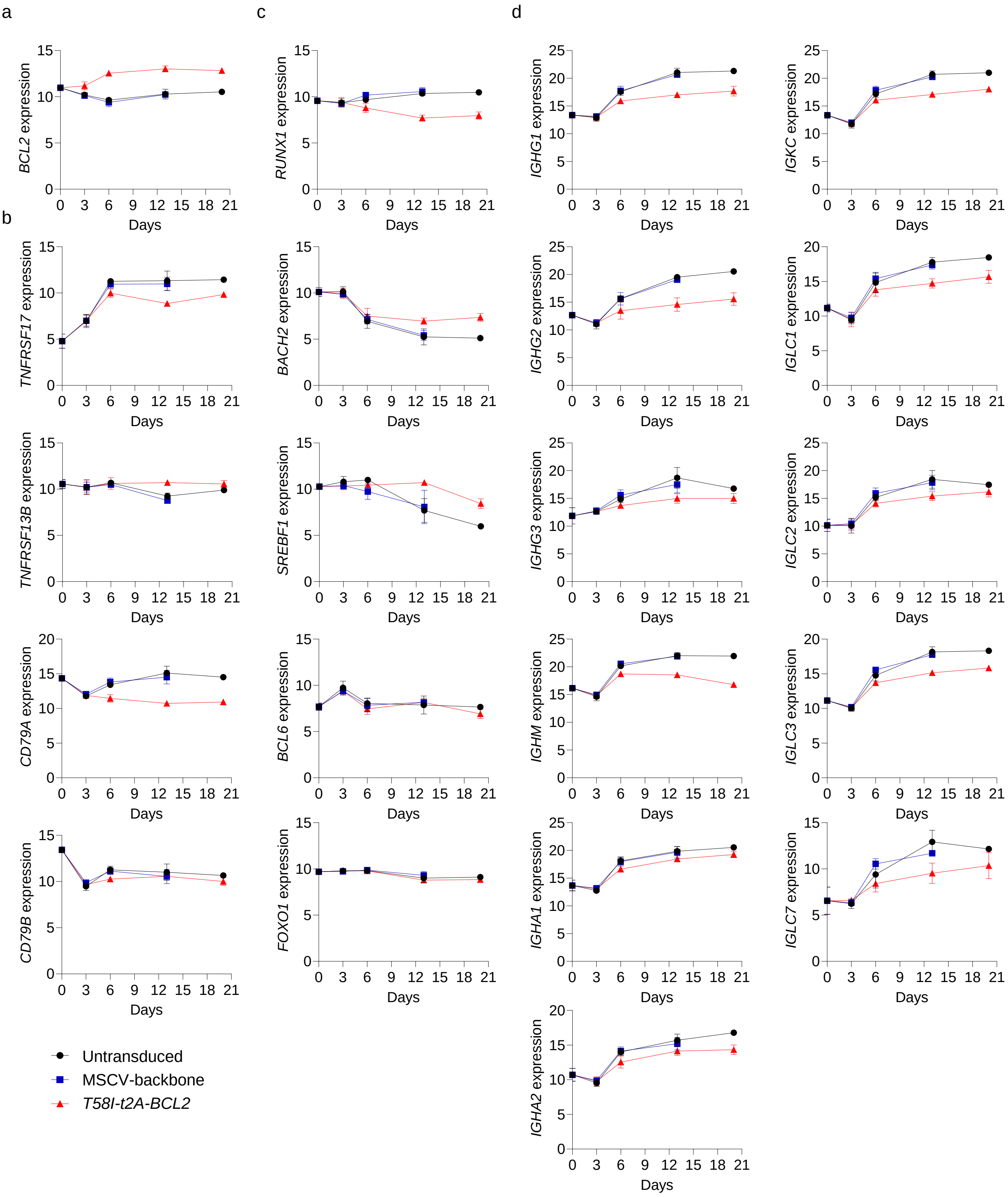


**Supplementary Figure 2. Gene expression changes upon MYC-BCL2 overexpression. (Accompanies Figure 2).** $Log_2$ normalised RNAseq expression values (y-axis) of selected genes indicated to the side of each graph and plotted per time point tested (x-axis) for the untransduced (black), MSCV-backbone (blue) and *T58I-t2A-BCL2* (red) conditions as indicated. Shown are expression levels of **a,** BCL2; **b,** surface proteins; **c,** transcription factors; and **d,** immunoglobulin heavy and light chain genes. Data are representative of two independent experiments with n=1-4 samples total depending on time point/condition.
