## Supplementary Figure 3 for "Enforced MYC expression selectively redirects transcriptional programs during human plasma cell differentiation"

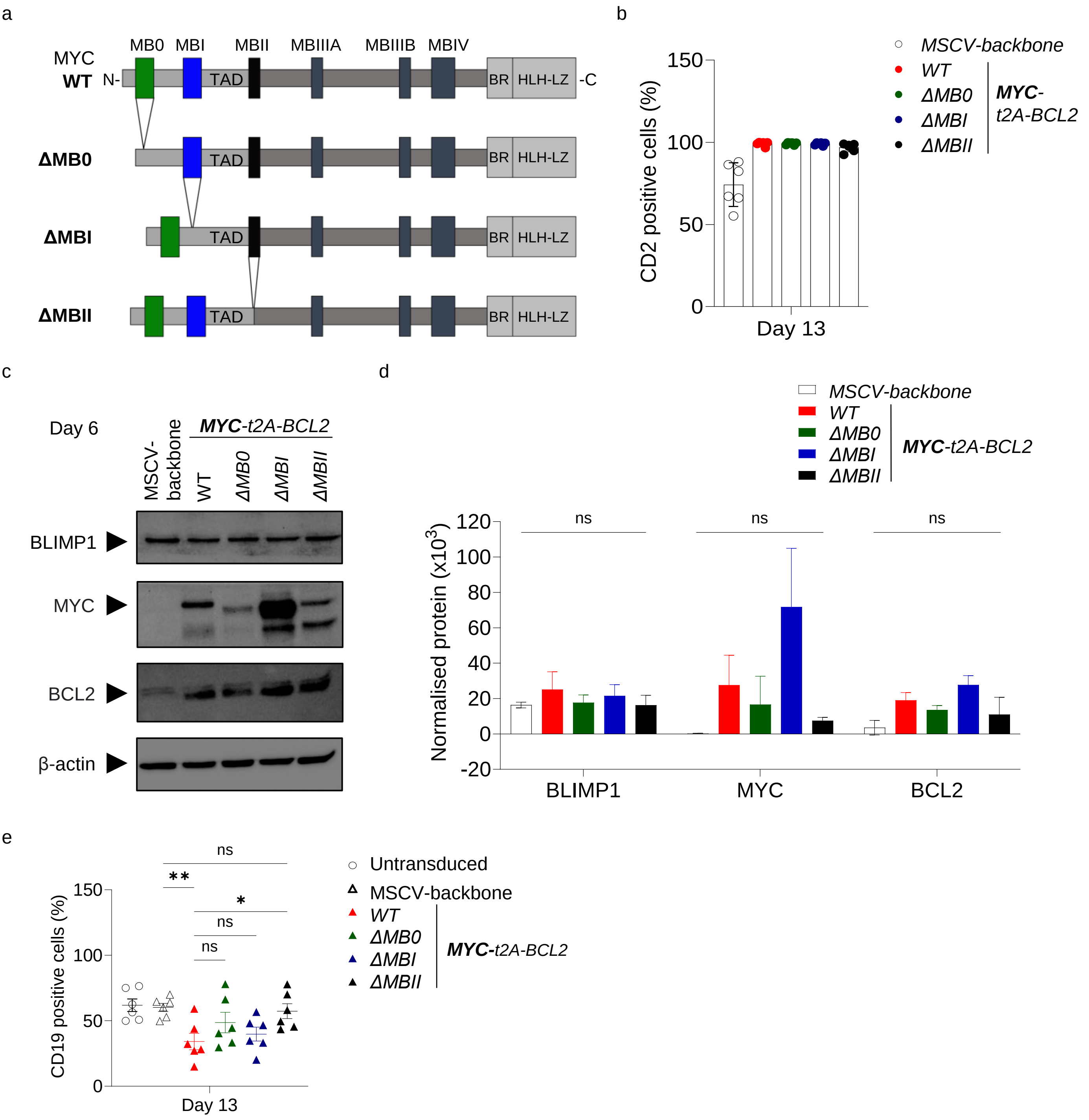


**Supplementary Figure 3. MYC TAD MB deletion mutants. (Accompanies Figure 4) a,** Graphical representation of MYC wild type versus the ΔMB0, ΔMBI, and ΔMBII mutants. **b,** Graph of flow cytometrically defined percentages of CD2 positive cells at day 13 for the indicated conditions. **c,** Western blot for BLIMP1, MYC, BCL2 and β-actin in total protein lysates of day 6 cells for conditions indicated above. **d,** Protein quantification normalized to β-actin loading control at day 6 for the indicated conditions. One-way ANOVA. **e,** Graph of the CD19 positive cell frequencies at day 13 from the *ΔMB0-*, *ΔMBI-*, and *ΔMBII-t2A-BCL2* flow cytometry data for the indicated conditions. Data shown for the conditions tested, apart from the untransduced, are pre-gated to $CD2^+$ populations. Data are representative of at least two independent experiments. Bars and error represent mean and standard deviation (SD); Unpaired two-tailed Student's *t-test.* ns, not significant; * $P < 0.05$; ** $P < 0.01$.
