## Supplementary Figure 4 for "Enforced MYC expression selectively redirects transcriptional programs during human plasma cell differentiation"

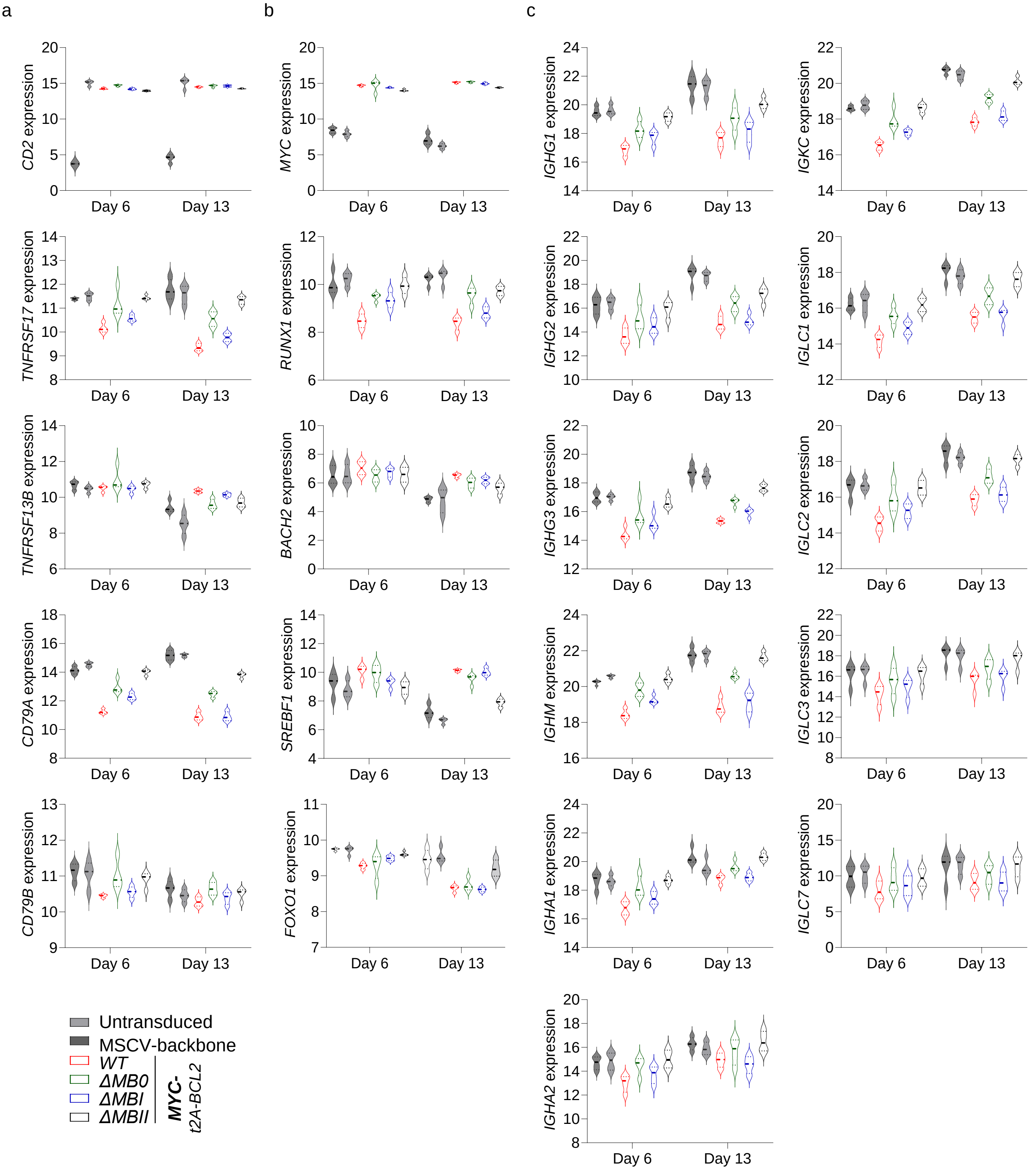


**Supplementary Figure 4. Gene expression changes for MYC MB deletions. (Accompanies Figure 5).** Violin plots of $log_2$ normalised RNAseq expression values (y-axis) of selected genes indicated to the side of each graph and plotted per time point tested (x-axis) for the conditions as indicated at bottom left of the figure. Shown are expression levels of **a,** surface proteins; **b,** transcription factors; and **c,** immunoglobulin heavy and light chain genes. Data are representative of two independent experiments with a total of n=4 samples per time point and condition.
