## Supplementary Figure 5 for "Enforced MYC expression selectively redirects transcriptional programs during human plasma cell differentiation"

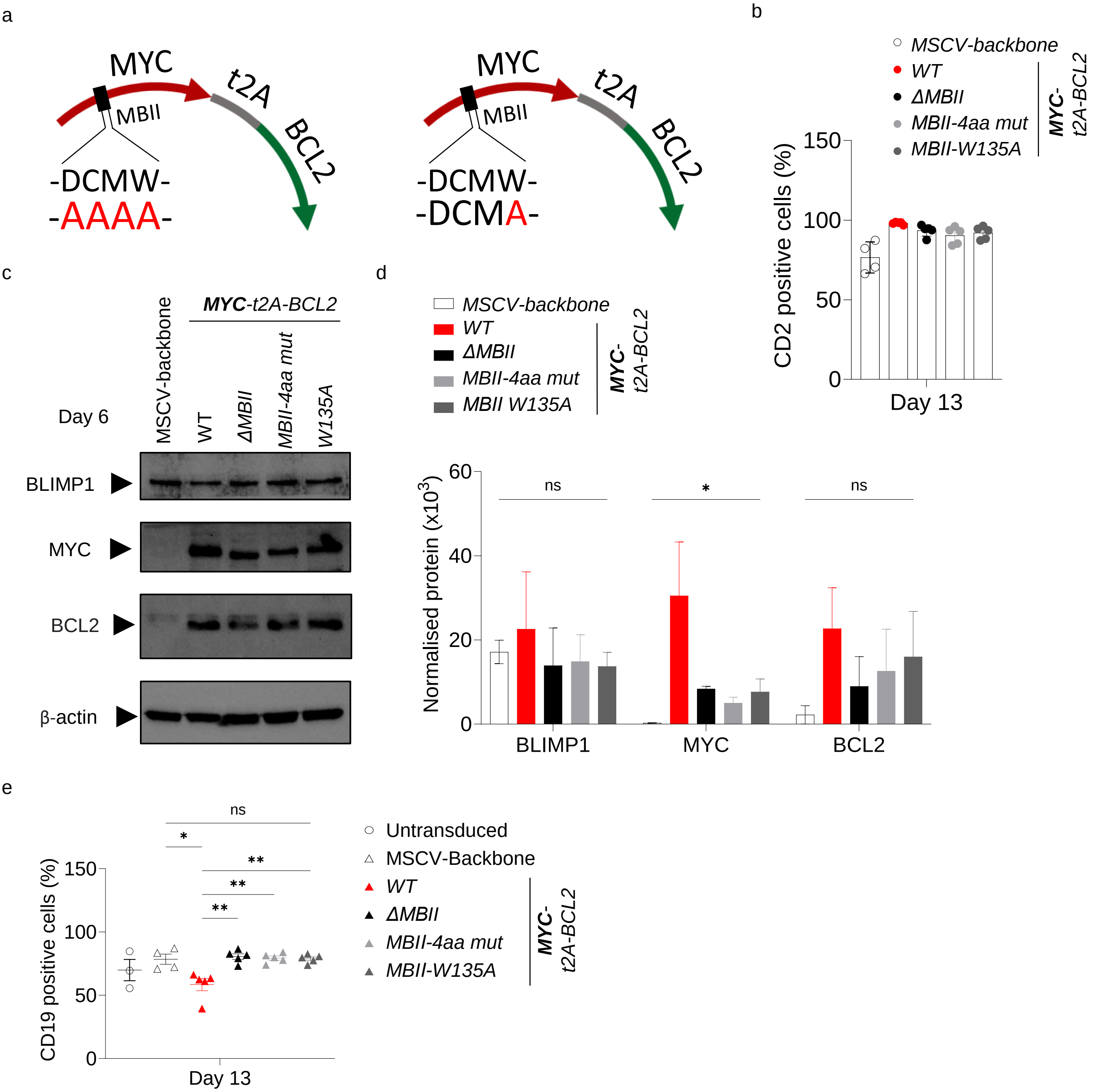


**Supplementary Figure 5. Details of MBII domain mutants of DCMW and W135. (Accompanies Figure 6) a,** Graphical representation of the MBII mutants, showing DCMW, motif mutated into four alanine residues (left) and the single mutation of W135A (right). **b,** Graph of flow cytometry percentages of CD2 positive cells at day 13 for the indicated conditions. **c,** Western blot of total protein lysates collected at day 6 showing BLIMP1, MYC, BCL2, and β-actin detection in the indicated conditions. **d,** Protein quantification normalized to β-actin at day 6 for the indicated conditions. Bars and error represent mean and standard deviation (SD). One-way ANOVA. **e,** Graph of flow cytometric CD19 positive cell frequencies at day 13 for the indicated conditions. Data shown for the conditions tested, apart from the untransduced, are pre-gated to $CD2^+$ populations. Data are representative of at least two independent experiments; Unpaired two-tailed Student's *t-test*. ns, not significant; * $P < 0.05$; ** $P < 0.01$.
