## Supplementary Figure 6 for "Enforced MYC expression selectively redirects transcriptional programs during human plasma cell differentiation"

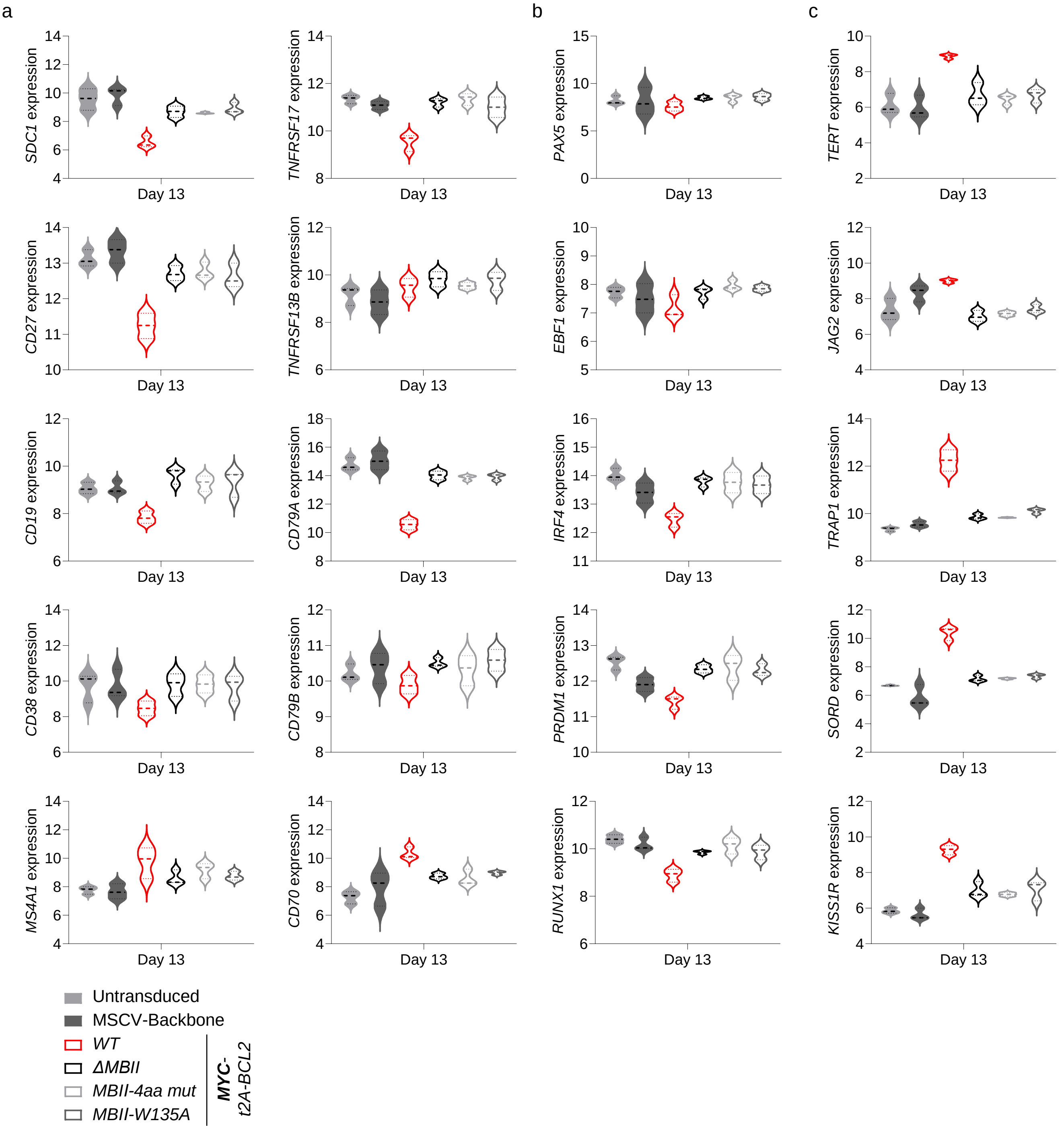


**Supplementary Figure 6. Gene expression changes for MBII mutant conditions. (Accompanies Figure 7).** Violin plots of $log_2$ normalised RNAseq expression values (y-axis) of selected genes indicated to the side of each graph at day 13 for the conditions as indicated at bottom left of the figure. Shown are expression levels of **a,** surface proteins; **b,** transcription factors; and **c,** MYC target genes. Data are representative of two independent experiments with a total of n=3 samples per time point and condition.
