## Supplementary Figure 7 for "Enforced MYC expression selectively redirects transcriptional programs during human plasma cell differentiation"

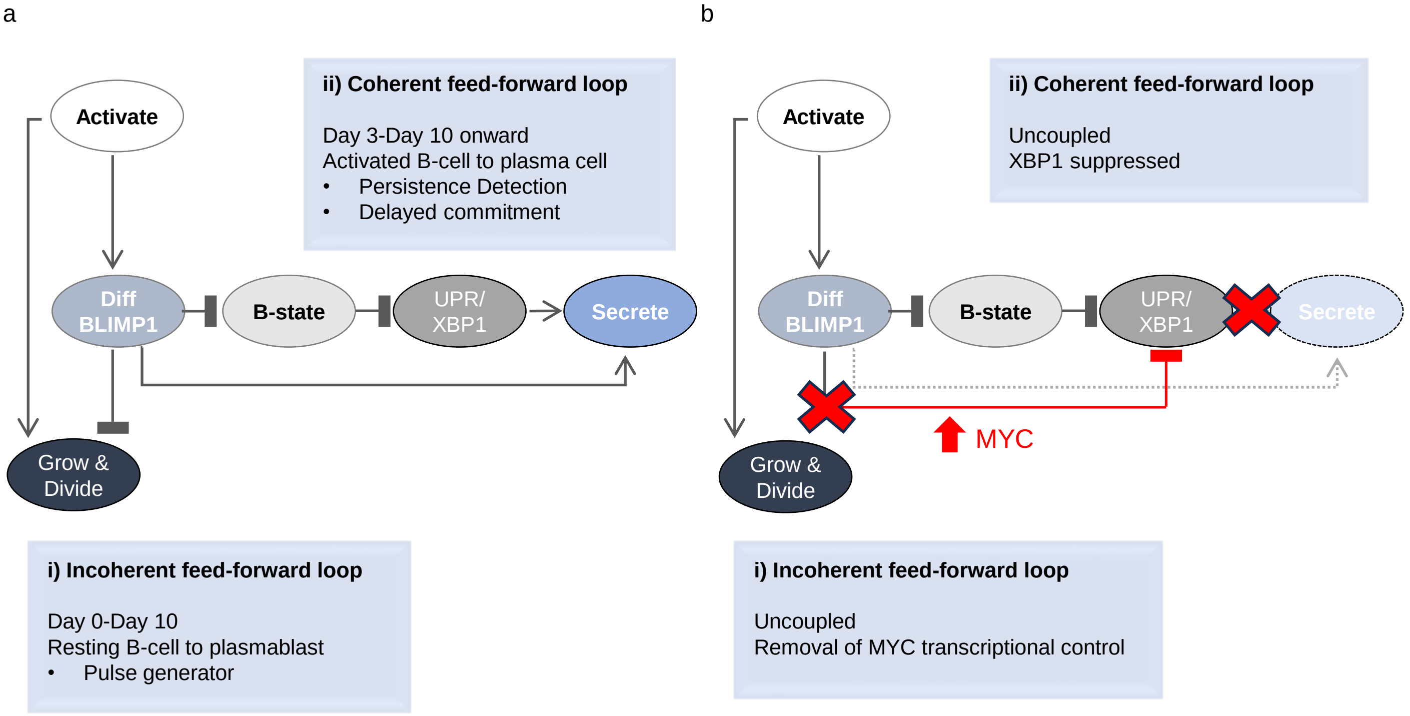


**Supplementary Figure 7. Graphical summary a,** During B-cell to plasma cell differentiation the arrangement of transcriptional regulation is consistent with coupled feedforward loops: i) The physiological MYC associated growth program is regulated by opposing inputs (incoherent feedforward loop). First it is controlled by positive action of input signals and then after a delay by the opposing negative action of the IRF4 and BLIMP1 associated differentiation (Diff) program. This enables a pulse of cell growth and division which is eventually curtailed; ii) transition to the secretory program is controlled by a coherent feedforward loop. In this arrangement BLIMP1 and XBP1, the key transcriptional regulator of the unfolded protein response of the ER (UPR), ultimately co-operate to support optimal secretory reprogramming. The delay in secretory commitment which is coupled with the burst of growth and proliferation may be explained by a need for repression of the B-cell state program (B-state) to allow XBP1 derepression (PAX5 has been implicated as a direct repressor of XBP1). **b**, Enforced expression of MYC from the activated B-cell state onward disrupts both feedforward loops: i) enforced MYC expression disrupts the early control over cell growth and division, likely by separating MYC expression from BLIMP1 mediated repression; ii) enforced MYC expression disrupts the coherent feedforward loop controlling transition to the secretory state. This is linked to repression of XBP1 despite suppression of the B-cell state. Control over the suppression of key transcription factors of the B-cell state is not impacted by MYC overexpression. Thus, distinct aspects of plasma cell differentiation are separable by feedforward loop uncoupling.
