## Supplementary material for "Enforced MYC expression selectively redirects transcriptional programs during human plasma cell differentiation": Methods

*Peripheral blood donors and cell lines*

Peripheral blood was obtained from leukocyte cones of healthy anonymous donors (NHSBT) or by blood collection from healthy volunteers following the appropriate consent requirements. HEK-293 cells were maintained in the culture using DMEM medium (41965039, Thermofisher Scientific) with 10% heat-inactivated fetal bovine serum (HIFBS) and 1:100 penicillin/streptomycin (15140122, Gibco) (DMEM CM). Irradiated murine fibroblasts transfected with human CD40L-L (CD40L-L stromal cells) were cultured in IMDM (31980-022, Gibco) with 10% HIFBS (IMDM CM). Cells were incubated at 37 °C with 5% CO_2_.

*Plasmids and cloning*

The retroviral construct *T58I-t2A-BCL2*, containing human *MYC* sequence, with T58I substitution, in combination with *BCL2* and a CD2 reporter*,* was kindly provided by the Daniel Hodson group as well as the packaging and envelop plasmids pHIT60 and GALV-MTR, respectively [1, 2]. A *WT-t2A-BCL2* construct containing the *MYC* wild type (WT) sequence in combination with *BCL2*, was also designed and synthesized commercially. To produce the TAD MBs deletion mutants, DNA fragments of the MB0, MBI, and MBII MYC domains corresponding to 5’- YDSVQPYFYCDEEENFY -3’, 5’- PSEDIWKKFELLPTPPLSP -3’ and 5’- IIIQDCMWSGFSAAAK -3’, respectively, were designed to be deleted from the *WT-t2A-BCL2* insert. The MYC TAD MBs deletion mutants as well as the mutation of MYC 132-135 amino acid sequence, DCMW, to alanine residues and the single substitution W135A were commercially synthesized with the *t2A-BCL2* sequence included in the insert and cloned into the pIRES2-EGFP vector (Clonotech). All the *MYC* mutants containing *t2A-BCL2* were subcloned into the MSCV-IRES-huCD2 plasmid kindly provided by the Daniel Hodson group. Plasmid propagation and successful ligations took place using NEB Stable Competent E. coli (C3040 NEB) for all the MSCV-based constructs and DH5a E. coli (18265-017, Invitrogen) for pIRES2-EGFP plasmids following manufacturer’s instructions, respectively. Diagnostic restriction enzyme digests verified all the constructs and additional Sanger sequencing validated the MYC 132-135 amino acids mutation into alanine and the W135A mutant. Plasmids were stored at -20 °C.

*Human memory B cell differentiation system*

Peripheral blood mononuclear cells (PBMCs) were isolated by Lymphoprep density gradient, 2000 rpm centrifugation for 20 minutes at room temperature (RT). PBMCs were washed with phosphate saline buffer (PBS), counted, and labeled appropriately, based on a human memory B cells isolation kit (130-093-546, Miltenyi Biotec), with B cell biotin antibody cocktail for 20 minutes at 4 °C. Magnetic isolation of total B cells was performed upon incubation for 20 minutes at 4 °C with anti-biotin beads. Following the previously established *in vitro* plasma cell differentiation method [3-5], memory B cells were isolated following magnetic labeling with anti-CD23 beads (130-094-510, Miltenyi Biotec) and co-cultured in a 24 well-plate format at 2x10^5^ cells/ml with 2x10^4^ cells/ml CD40L-L stromal cells in IMDM CM containing IL-2 (40 U/ml), IL-21 (100 ng/ml) and F(ab’)2 fragments goat anti-human IgG, IgM and IgA (20 µg/ml) (109-006-064, Jackson Immunoresearch). At day 3 of B cell differentiation *in vitro*, activated B cells were seeded without CD40L-L stromal cells in a new 24 well-plate at 0.33x10^5^ cells/ml for the *T58I-t2A-BCL2*, *WT-t2A-BCL2* and *ΔΜΒΙ-t2A-BCL2* conditions and at 1x10^5^ cells/ml for the rest of the conditions, in IMDM CM containing IL-2 (20 U/ml), IL-21 (50 ng/ml) and supplements (lipid mixture 1; chemically defined (200X) and MEM amino acids solution (50X)). At day 6, the *T58I-t2A-BCL2*, *WT-t2A-BCL2* and *ΔΜΒΙ-t2A-BCL2* cells were re-seeded at 0.66x10^5^ cells/ml, and the rest of the conditions at 2x10^6^ cells/ml in IMDM CM containing APRIL (100 ng/ml), IL-21 (10 ng/ml), IL-6 (10 ng/ml) and supplements at 1 ml final volume per well. At day 9, the *T58I-t2A-BCL2*, *WT-t2A-BCL2* and *ΔΜΒΙ-t2A-BCL2* cells were splitted following a 1:2 ratio, and appropriate volume of the day 6 complete medium was added to all the conditions aiming at a final volume of 2 ml per well. At day 13 onwards, the *T58I-t2A-BCL2*, *WT-t2A-BCL2* and *ΔΜΒΙ-t2A-BCL2* cells were re-seeded at 0.33x10^5^ cells/ml, and the rest of the conditions at 1x10^6^ cells/ml in IMDM CM containing APRIL (100 ng/ml), IL-6 (10 ng/ml) and supplements at 2 ml final volume per well.

*Retroviral production and viral stock validation*

HEK-293 cells seeded in 10 cm Petri dishes 24 hours in advance were transfected with 1 ml Opti-MEM (31985062, Invitrogen) mixed with 18 µl Transit-293T (MIR 2700, Mirus) transfection reagent containing 1 µg pHIT60 packaging, 1 µg GALV-MTR envelop and 4 µg retroviral constructs, as previously described [1, 2]. The virus was collected and filtered after 48 hours of incubation at 37 °C with 5% CO_2_ and either used fresh for transductions or aliquoted at 1 ml and stored at -80 °C. To validate the frozen viral stocks HEK-293 cells were seeded in 6 well plates 24 hours in advance and transduced with 1 ml frozen virus per well mixed with 10 µg/ml polybrene (sc134220, INSIGHT biotechnology). A spinfection step at 2500 rpm for 60 minutes at 30 °C was used to augment retroviral infection and the medium was replaced with fresh DMEM CM. CD2 staining, and flow cytometry assessment were conducted 72 hours post-transduction.

*Retroviral transduction of human memory B cells*

Human memory B cells co-cultured with CD40L-L were centrifuged at 1400 rpm for 4 minutes at RT. Subsequently, 80% of the growth medium was aspirated and the co-cultured activated memory B cells were transduced with 1 ml fresh or frozen retrovirus mixed with 25 µM HEPES (15630-056, Thermofisher Scientific) and 10 µg/ml polybrene (sc134220, INSIGHT biotechnology). A spinfection step at 2700 rpm for 90 minutes at 32 °C was used to augment retroviral infection and 70% of the medium was replaced with fresh IMDM CM containing IL-2 (20 U/ml), and IL-21 (50 ng/ml) [2-5].

*Flow cytometry*

Cells were washed with RT PBS and centrifuged at 1500 rpm for 5 minutes at RT. Live/dead fixable viability stain 780 nm (565388, BD Biosciences) was diluted 1:1000 in RT PBS, 500 µl were shared per sample and incubated for 15 minutes at RT. Cells were washed with 2 ml FACs buffer (PBS + 0.5% HIFBS) and centrifuged at 1500 rpm for 5 minutes at RT. 25 µl blocking buffer containing hIgG molecules (I2511-10MG, Sigma) and natal mouse serum in FACs buffer were added, and samples were incubated for 15 minutes at RT. 10 µl antibodies master mix was added followed by 20 minutes at RT incubation. Stained cells were washed with 2 ml FACs buffer and centrifuged at 1500 rpm for 5 minutes at RT. Cells were fixed with 150 µl 2% paraformaldehyde and stored at 4 °C for flow cytometry analysis. Intracellular staining was performed by permeabilizing the cells in saponin-based permeabilization buffer for 20 minutes and staining was followed for 1,5 hours at 4 °C. Antibodies used were: CD19-PE (130-113-169, Miltenyi), CD20-ef450 (48-0209-42, Invitrogen), CD20-BV421 (562873, BD Biosciences), CD27-FITC (555440, BD Pharmingen), CD38-PE-Cy7 (335825, BD Biosciences), CD138-APC (130-117-395, Miltenyi), CD2-BUV395 (563820, BD Biosciences), Ki67-Alexa fluor 488nm (558616, BD Biosciences). CountBright beads (C36950, Invitrogen) were used for absolute cell number analysis in combination with trypan blue-based haemocytometer counts. Flow cytometry data were collected using Cytoflex S and Cytoflex LX analysers (Beckman Coulter). Flow cytometry analysis was conducted using FlowJo software v.10.7.2 and v.10.8.1 and GraphPad Prism 10 software.

*Proliferation assays*

5-ethynyl-2'-deoxyuridine (EdU) 1 hour pulse assay took place for the day 21 and 31 *in vitro* differentiated untransduced and *T58I-t2A-BCL2* cells. EdU incorporation was detected using the Click-iT Plus EdU Alexa fluor 647 kit (C10635, Invitrogen) based on the provided protocol and followed by Ki67 intracellular staining as described above. Cellular proliferation was assessed by flow cytometry.

*Western blotting*

Cells were washed with sterile PBS and protein was extracted with 30 µl RIPA buffer. Lysed cells were kept on ice for 15 minutes and centrifuged at 12,000 rpm for 30 minutes at 4 °C following supernatant collection. BCA assay (AR0146, Boster or 23250, Thermo Scientific) was performed for protein quantification according to the manufacturer’s instructions. Normalised lysates were loaded in SDS-PAGE followed by wet-western blotting for the detection of c-MYC (D3N8F rabbit, 1:1000 13987S, Cell Signalling Technology), BLIMP1 (PRDM1a form mouse, 1:1000), BCL2 (2870S, Cell Signalling Technology, 1:1000), BCL2 (2872, Cell Signalling Technology, 1:1000) and β-actin (A1978-200UL mouse, 1:10,000). Horseradish peroxidase (HRP) conjugated rabbit or mouse secondary antibodies were used at 1:10,000. Membrane development was performed using enhanced chemiluminescent HRP substrate (34580, Thermo Fisher Scientific) incubation and images were taken with ChemiDoc MP Imaging System (BioRad). Membranes were stripped using stripping buffer (46430, Thermo Fisher Scientific) for 10 minutes at RT.

*Enzyme-Linked Immunosorbent Assay (ELISA)*

Human IgG (A80-104A, Bethyl) and human IgM (A80-100A, Bethyl) ELISA quantification sets were used to the manufacturer’s instructions for total IgG and IgM antibody secretion detection respectively. Standard curves were generated from readings of absorbance at 450 nm with a Cytation 5 imaging plate reader (BioTek). Analysis was conducted using MyAssays Ltd online data analysis tool and GraphPad Prism 10 software.

*RNA extraction*

Cells were counted and lysed with 800 µl to 1 ml TRIzol (15596026, ambion) reagent, incubated for 10 minutes at RT and stored at -80 °C. Chloroform (C2432-26ML, Honeywell) was added at 160 µl or 200 µl respectively, to defrosted samples followed by vigorous shake for 15 seconds and 3 minutes at RT incubation. A centrifugation step at 11,000 rpm for 15 minutes at 4 °C was followed by collection of the aqueous phase. 400 µl isopropanol and 10 µl glycogen (AM9510, Invitrogen) were added and the mixture was incubated for 10 minutes at RT. Samples were centrifuged at 11,000 rpm for 10 minutes at 4 °C and the pellet was washed three times with 75% ethanol followed by a centrifugation step at 9,000 rpm for 5 minutes at 4 °C. RNA pellets were air dried for 10 minutes and dissolved in 30 µl RNase free water. RNA was incubated at 55 °C for 10 minutes, treated with DNases for genomic DNA removal (AM1906, Invitrogen) and stored at -20 °C.

*RNA-seq analysis*

RNA-seq was conducted on a Novoseq6000 platform (Illumina), using 150-bp paired-end sequencing. The fastq files were assessed for initial quality using FastQC v0.11.8, trimmed for adapter sequences using TrimGalore v0.6.10 and aligned to GRCh38.p13/hg38 with STAR aligner (v2.6.0c) [6]. Transcripts were re-annotated with the MyGene.info API using all available references and any ambiguous mappings manually assigned. Transcript abundance was estimated in RSEM v1.3.1 and imported into R v4.1.2 with txImport v1.22.0 and then processed using DESeq2 v1.34.0 [7-10]. Software DESeq2 determined differential gene expression (DEG) between every contrast and a total DEG carried out with a likelihood ratio test (LRT), quality visualised using MA plots and shrinkage of log fold estimated using the apeglm method (Supplemental Table 1) [11].

*RNA-seq network*

Parsimonious Gene Correlation Network Analysis (PGCNA) approach was utilised, for details and validation of the PGCNA approach see our other work [12]. The transcripts differentially expressed between any contrast or across the timeseries data (DESeq2 FDR < 0.01) were retained for PGCNA analysis (n=14,360 genes) giving a 14,360 x 48 matrix. This was used for a PGCNA2 analysis (-n 1000, -b 100) giving a network with multiple modules. The median expression per condition/time was visualised as Z-scores mapped onto the network. For each gene in the network a strength (edge-weight x degree) was calculated and used to select the top 10 genes per module. These were converted to Module Expression Values by taking their median Z-scores (across samples) and visualised as a hierarchically clustered heatmap.

*Enrichment analysis*

The gene signature enrichment (GSE) was assessed using a hypergeometric test, in which the draw is the gene list genes, the successes are the signature genes, and the population is the genes present on the platform. The resultant p-values are then adjusted for multiple testing using Benjamini and Hochberg correction. For the PGCNA networks GSE analyses the genes per module were compared against the 43,572-signature database (background: 14,360 genes in network). Only signatures that contain at least 3 genes in background set were retained.

*Data processing and availability*

All RNA-seq data analyses were undertaken on ARC4, part of the High-Performance Computing facilities at the University of Leeds, UK. Interactive networks and all meta-data are available at <https://matthewcare.wixsite.com/pgcna/myc-trd>. PGCNA python scripts are available at <https://github.com/medmaca/PGCNA>.

The primary datasets are available at the Gene Expression Omnibus [GSE262809](https://eur03.safelinks.protection.outlook.com/?url=https%3A%2F%2Fwww.ncbi.nlm.nih.gov%2Fgeo%2Fquery%2Facc.cgi%3Facc%3DGSE262809&data=05%7C02%7CR.Tooze%40leeds.ac.uk%7C222f201f0b184e72be2a08dc53bb0522%7Cbdeaeda8c81d45ce863e5232a535b7cb%7C0%7C0%7C638477309914300929%7CUnknown%7CTWFpbGZsb3d8eyJWIjoiMC4wLjAwMDAiLCJQIjoiV2luMzIiLCJBTiI6Ik1haWwiLCJXVCI6Mn0%3D%7C0%7C%7C%7C&sdata=pgGZrYVBI6xDt%2FBDpfGzAsYaLuRY0ooKzA58KQOtJTQ%3D&reserved=0).

*Statistical analysis*

Statistical tests were performed using one-way ANOVA or unpaired two-tailed Student’s *t* test with GraphPad Prism 10 software.

*Ethical approval*

Approval for this study was provided by UK National Research Ethics Service via the Leeds East Research Ethics Committee, approval reference: 07/Q1206/47, IRAS reference 187050.

**Methods References**
